# Hydrophobic Patch Spacing Produces Nonmonotonic Compaction in Intrinsically Disordered Proteins

**DOI:** 10.64898/2026.09.02.748969

**Authors:** Henry Silvernail, Wangfei Yang, Wenwei Zheng

## Abstract

The conformational ensembles of intrinsically disordered proteins (IDPs) are encoded by the distribution of physicochemical interactions along their sequences. Although hydropathy-based descriptors capture average chain dimensions across diverse IDPs, the consequences of spacing localized hydrophobic patches remain poorly understood. Coarse-grained simulations of fixed-composition FUS-derived sequence variants reveal maximal compaction at an intermediate patch spacing. Analysis of simplified model peptides identifies this nonmonotonic behavior as one of three spacing responses: monotonic expansion, nonmonotonic compaction, and monotonic compaction. Their occurrence depends on interaction strength, effective interaction length scale, and patch architecture, with nonmonotonicity emerging only when hydrophobic attractions are sufficiently strong. A conformational-class decomposition resolves these responses into weighted patch-contact and patch-noncontact contributions to the ensemble-averaged chain dimensions. In attractive regimes, the contact contribution decreases with spacing while the noncontact contribution increases, and the changing balance between these opposing effects produces maximal compaction at an intermediate spacing. In the steric-dominated regime, separating the patches instead compacts the dominant noncontact conformations by relieving steric frustration. These steric-and attraction-dominated limits show that similar spacing responses can arise from distinct microscopic mechanisms. These findings establish a unified framework for understanding how interaction regime and hydrophobic patch spacing jointly shape IDP conformational ensembles.

## Introduction

Intrinsically disordered proteins or regions (hereafter IDPs) perform essential regulatory and signaling functions and are implicated in numerous pathological processes despite lacking well-defined tertiary structures.^1–4^ Their structural behavior is instead described by dynamic conformational ensembles that mediate molecular recognition, biomolecular condensation, and aberrant aggregation.^5–9^ These ensembles are encoded not only by amino acid composition but also by the linear arrangement of physicochemical interactions along the sequence.^10^ Establishing how one-dimensional sequence organization maps onto threedimensional conformational ensembles is therefore central to understanding and engineering IDPs and IDP-driven condensates for specific functions.^11,12^ A major unresolved aspect of this problem is how sequence separation acquires different physical meanings for different classes of molecular interactions.

Recent proteome-scale analyses have shown that nonrandom residue-pair patterning defines IDP molecular grammars associated with distinct biological functions and cellular localization.^13^ Charge patterning provides one of the clearest mechanistic examples of how such sequence organization controls conformational ensembles. Early work established that net charge and the fractions of charged residues strongly influence IDP conformations, ^3,14^ while subsequent studies showed that rearranging charged residues at fixed composition can also produce substantial changes in chain dimensions. These observations motivated charge-patterning descriptors such as *κ* ^15^ and Sequence Charge Decoration (SCD).^16,17^ Electrostatic interactions are signed, producing attraction between opposite charges and repulsion between like charges, and act over comparatively long spatial distances before being screened by solution ions.^18^ Consistent with this physics, SCD weighs each charge pair by the square root of its sequence separation, allowing pairs separated by larger contour distances to exert greater influence on global chain dimensions. This sequence-separation dependence emerges from analytical polymer-theory treatment of electrostatic interactions and has provided a physical foundation for understanding charge patterning in IDPs.^16,17,19,20^

Hydrophobic interactions coexist with electrostatic interactions in IDPs, and their relative contributions to sequence-dependent assembly can vary substantially with sequence context.^21^ Unlike signed, comparatively long-ranged electrostatic interactions, hydrophobic interactions are predominantly attractive and short-ranged, giving hydrophobic sequence organization a distinct physical basis. Although individual residue-level attractions may be modest, clustering hydrophobic residues into patches can generate strong effective associations through multivalent contacts. Recent studies have established localized hydrophobic and aromatic clusters as important sequence elements regulating IDP conformational ensembles, phase behavior, and functions.^22–26^ Hydropathy-based descriptors, including Sequence Hydropathy Decoration (SHD)^27^ and local hydrophobic clustering (HpC), ^28^ have substantially improved predictions of conformational properties across diverse IDP sequences and have been incorporated into a sequence design framework.^29^ SHD was developed as a hydropathy analogue of SCD and weights hydrophobic residue pairs inversely with their separation along the sequence, thereby assigning progressively smaller contributions to residues separated farther along the sequence. The contrasting sequence-separation exponents in SCD and SHD should not be interpreted as direct representations of the spatial ranges of the underlying interactions. Rather, they indicate that sequence separation couples differently to signed, comparatively long-ranged electrostatic interactions and short-ranged, contact-mediated hydrophobic interactions. Although the inverse weighting in SHD successfully captures average conformational trends, whether hydrophobic spacing can be represented by a single monotonic weighting across different interaction regimes and patch architectures remains unknown. For localized hydrophobic patches, increasing sequence separation may simultaneously reduce the probability of patch contact and increase the degree of chain compaction when contact occurs. How these competing effects regulate chain compaction has not been systematically established. Residue-level coarse-grained (CG) models provide a useful framework for isolating such sequence-dependent effects under controlled interaction conditions.^30–33^

Here, we use CG molecular dynamics simulations^33^ to determine how hydrophobic patch spacing governs IDP conformational ensembles. We examine composition-preserving sequence variants derived from the low-complexity domain of Fused in Sarcoma (hereafter referred to as FUS),^34^ together with simplified model peptides spanning different interaction regimes and sequence architectures. We show that hydrophobic patch spacing can produce distinct monotonic and nonmonotonic conformational responses through competition between the decreasing probability of patch contact and the greater chain compaction produced when patches farther apart in sequence form contacts. Extending this analysis to repulsive patterning further demonstrates that the conformational effect of sequence separation depends on the interaction regime, providing a physical framework for extending sequence-based descriptions of IDP conformational ensembles.

## Results and Discussion

### Hydrophobic Patch Spacing Produces Nonmonotonic Compaction

To examine how hydrophobic sequence organization influences IDP chain dimensions, we first investigated sequence variants derived from the FUS low-complexity domain using coarse-grained (CG) molecular dynamics simulations. Tyrosine is abundant in the FUS low-complexity domain and acts as a major associative residue.^34–38^ It is also among the residues with the highest CG-model hydropathy in this sequence.^33,39^ We therefore reorganized the tyrosine residues into two contiguous patches while preserving the order of the remaining residues (Fig. 1A). Patch spacing, *s*, was defined as the number of residues separating the nearest edges of the two hydrophobic patches, with *s*=0 corresponding to adjacent patches. Across the sequence variants, patch spacing was systematically varied without changing the overall amino acid composition, thereby isolating the effect of sequence organization from composition. A representative simulated conformation is shown in Fig. 1B.

**Figure 1:**
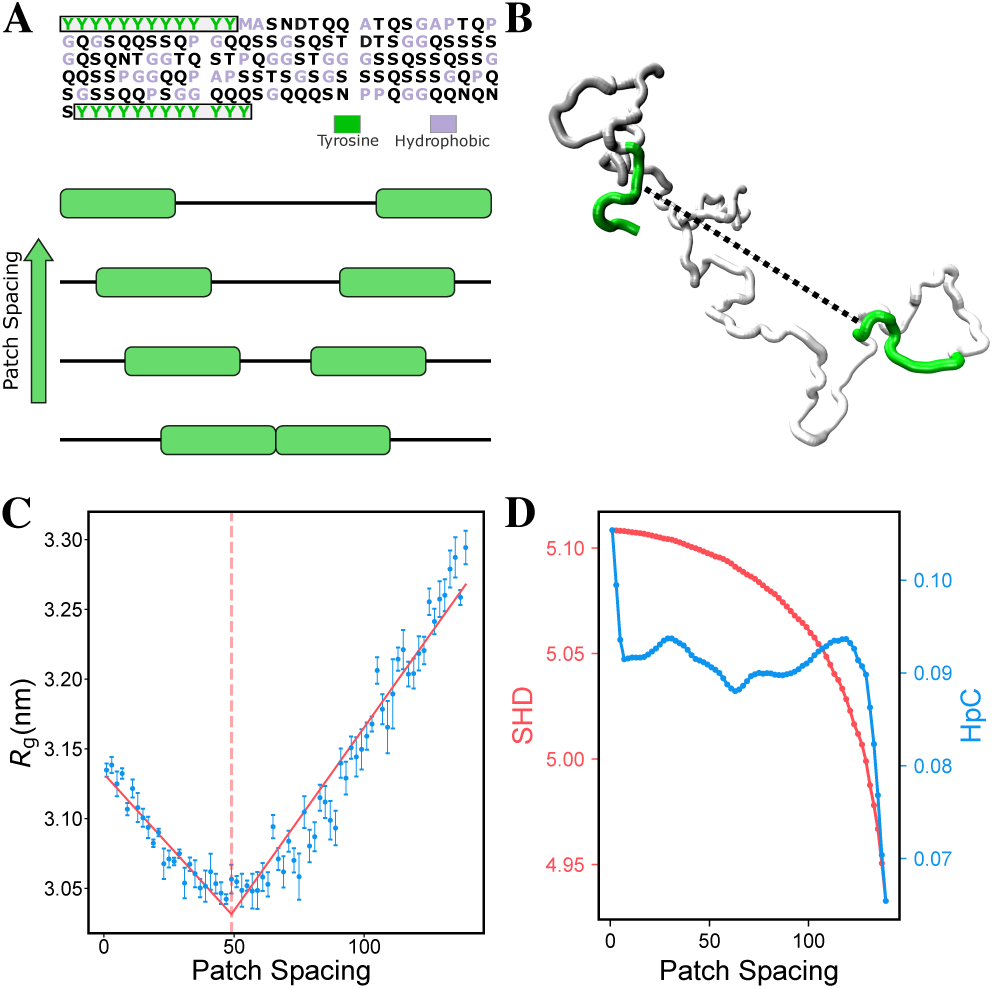
Hydrophobic patch spacing produces nonmonotonic chain compaction in FUS-derived sequence variants. (A) Sequence and schematic representation of composition-preserving variants derived from the FUS low-complexity domain. Tyrosine residues (green) are arranged into two contiguous hydrophobic patches separated by a variable number of intervening residues, while other hydrophobic residues are shown in light purple. (B) Representative simulated conformation showing the two hydrophobic patches (green). (C) Radius of gyration (*R_g_*) as a function of hydrophobic patch spacing. Red lines represent piecewise linear fits, and vertical dashed lines indicate the corresponding optimal hydrophobic spacing (see Methods). (D) Corresponding SHD (red) and HpC (blue) values as a function of patch spacing.

Patch spacing did not produce a simple monotonic change in chain dimensions. Instead, the radius of gyration (*R_g_*) exhibited a clear nonmonotonic dependence on patch spacing (Fig. 1C). For the 12-residue patches, the chain initially became more compact as patch spacing increased, reached maximal compaction at an intermediate spacing, and subsequently expanded at larger spacings. The same qualitative response was observed for patch lengths of 8 and 10 residues (Fig. S1), indicating that the nonmonotonic behavior is not restricted to a single patch size.

The hydropathy-based sequence descriptors examined here did not reproduce this non-monotonic response. As shown in Fig. 1D, Sequence Hydropathy Decoration (SHD)^27^ changes monotonically with patch spacing because it assigns progressively smaller weights to hydrophobic residue pairs separated farther along the sequence and therefore predicts monotonic chain expansion. In contrast, local hydrophobic clustering (HpC)^28^ is comparatively insensitive across most intermediate patch spacings because it primarily captures local clustering rather than the separation between distant patches. Neither descriptor identifies the spacing of maximal compaction, indicating that pairwise hydropathy weighting by sequence separation and local clustering alone do not capture the collective conformational response of extended hydrophobic patches.

These results show that hydrophobic patch spacing can regulate IDP chain compaction in a manner not anticipated by current sequence descriptors. They raise two central questions: under what physical conditions does nonmonotonic compaction emerge, and what molecular mechanism produces it? To address these questions, we next turn to simplified model polymers in which interaction strength, effective interaction length scale, and patch architecture can be varied systematically.

### Interaction Strength Determines the Emergence of Nonmonotonic Compaction

To isolate the role of hydrophobic interactions from sequence-specific effects, we constructed a simplified 150-residue peptide containing two hydrophobic patches separated by a variable number of intervening residues. The interaction strengths of the patch and nonpatch residues, quantified in the simulation model by hydropathy parameters *λ*_1_ and *λ*_2_, respectively, were varied independently while maintaining a constant patch length of 12 residues.

Systematic variation of *λ*_1_ and *λ*_2_ revealed qualitatively different responses to hydrophobic patch spacing (Fig. 2A). At weak interaction strengths, *R_g_* increased monotonically with patch spacing. In contrast, a clear minimum in *R_g_* emerged when both the patch and nonpatch attractions were sufficiently strong. Within this nonmonotonic regime, the spacing of maximal compaction shifted toward larger values as the interaction strengths increased. Thus, nonmonotonic compaction is not an intrinsic consequence of hydrophobic patch spacing but emerges only within a sufficiently attractive region of the interaction parameter space.

**Figure 2:**
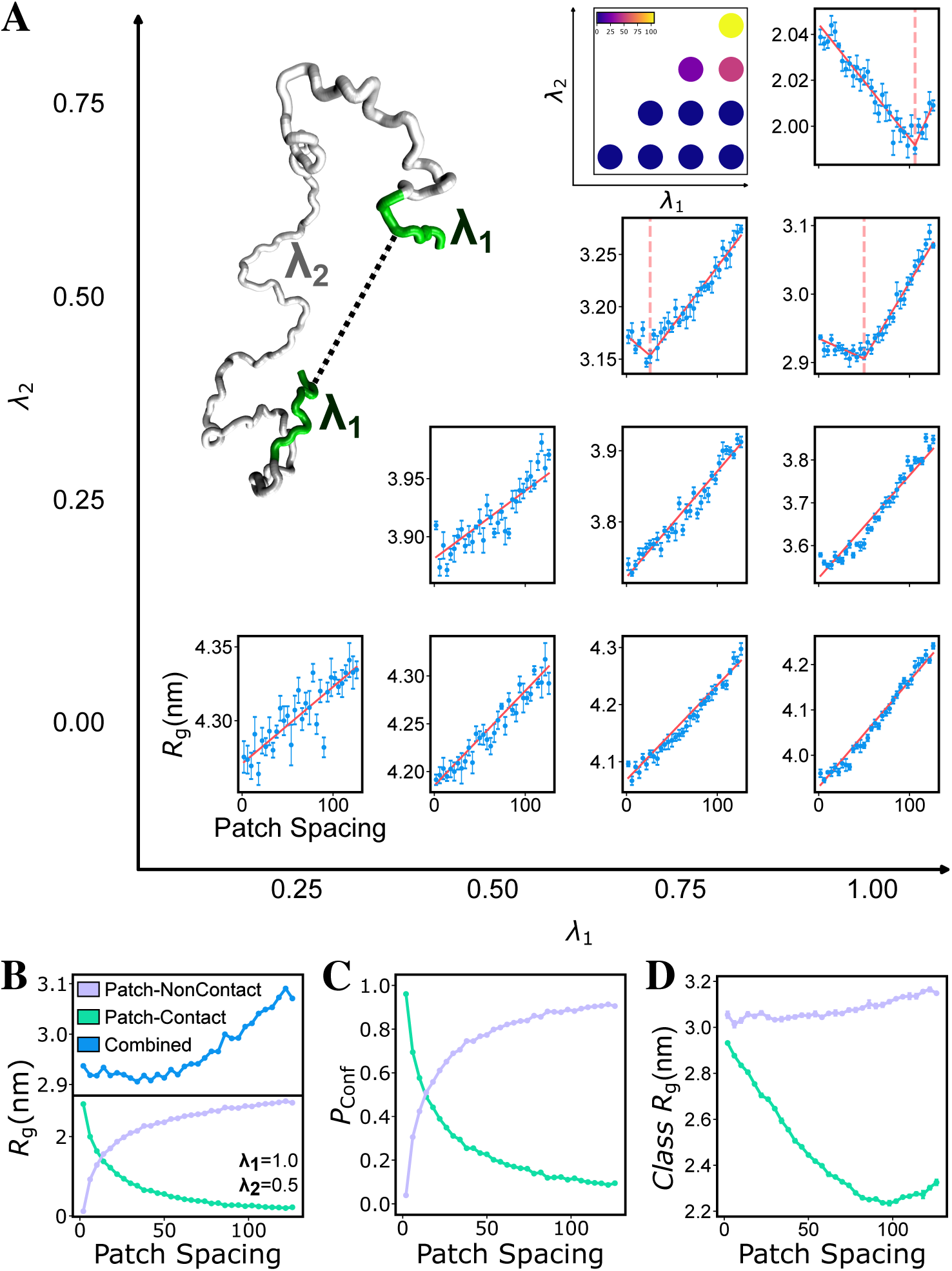
Interaction strength determines the emergence of nonmonotonic compaction. (A) *R_g_* as a function of hydrophobic patch spacing for 150-residue model peptides containing two 12-residue hydrophobic patches, shown for different combinations of the patch (hydropathy parameter *λ*_1_) and nonpatch (*λ*_2_) interaction strengths. Red lines represent piecewise linear fits used to determine the spacing of maximal compaction (see Methods). The parameter-space inset summarizes the fitted optimal patch spacing, with zero indicating monotonic expansion for which no interior minimum was assigned. A representative conformation is shown with the hydrophobic patches in green and the nonpatch residues in gray. (B) Weighted *R_g_* contributions of the patch-contact and patch-noncontact conformational classes, calculated as *P*_contact_*R_g,_*_contact_ and *P*_noncontact_*R_g,_*_noncontact_, respectively. Their sum, corresponding to the ensemble-averaged *R_g_*, is shown in the upper panel. (C) *P*_contact_ and *P*_noncontact_. (D) *R_g,_*_contact_ and *R_g,_*_noncontact_. Panels B–D show *λ*_1_ = 1.0 and *λ*_2_ = 0.5.

To determine the molecular origin of nonmonotonic compaction in the strongly attractive regime, we partitioned each conformational ensemble into patch-contact and patch-noncontact classes according to whether any residue pair belonging to different hydrophobic patches was within a certain contact-distance cutoff (see Methods). For each trajectory, we calculated both the population fraction of each conformational class (Fig. 2C) and its average *R_g_* (Fig. 2D). The ensemble-averaged *R_g_* can therefore be decomposed as

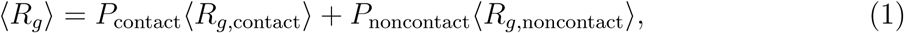

where *P*_contact_ and *P*_noncontact_ denote the population fractions of the patch-contact and patch-noncontact classes, respectively, and *P*_noncontact_ = 1 − *P*_contact_.

The two terms in Eq. 1 represent the weighted contributions of the patch-contact and patch-noncontact conformational classes to the ensemble-averaged *R_g_*. These contributions are shown in Fig. 2B for the representative condition *λ*_1_ = 1.0 and *λ*_2_ = 0.5. The weighted patch-contact contribution decreased monotonically with increasing patch spacing, whereas the weighted patch-noncontact contribution increased monotonically. At smaller spacings, the decrease in the patch-contact contribution exceeded the increase in the patch-noncontact contribution, causing the ensemble-averaged *R_g_* to decrease. At larger spacings, the patch-contact contribution began to level off while the patch-noncontact contribution continued to increase, causing the ensemble-averaged *R_g_* to turn upward. The nonmonotonic response therefore arises from the changing balance between the two opposing weighted contributions.

We further examined the population and average *R_g_* of each conformational class. The probability of patch contact decreased continuously with increasing patch spacing (Fig. 2C), with a corresponding increase in the patch-noncontact population. At the same time, *R_g,_*_contact_ decreased substantially over the initial and intermediate spacing range, whereas *R_g,_*_noncontact_ increased only modestly (Fig. 2D). These changes produce competing effects on the ensemble-averaged chain dimensions. The simultaneous decreases in *P*_contact_ and *R_g,_*_contact_ produce the monotonic decrease in the weighted patch-contact contribution. Conversely, the increase in *P*_noncontact_, reinforced by the modest increase in *R_g,_*_noncontact_, produces the increasing patch-noncontact contribution. At smaller spacings, the decrease in the patch-contact contribution dominates, causing the ensemble-averaged *R_g_* to decrease. At larger spacings, the increase in the patch-noncontact contribution dominates, producing the subsequent expansion.

The same conformational-class decomposition across the attractive interaction regimes reveals how interaction strength regulates the balance between these contributions (Fig. S2). The qualitative spacing dependence of *R_g,_*_contact_ remains similar across different interaction strengths, with the patch-contact conformations becoming more compact over the initial and intermediate spacing range (Fig. S3). In contrast, the magnitude and persistence of *P*_contact_ depend strongly on both *λ*_1_ and *λ*_2_ (Fig. S4). Within the intermediate and strong attraction regimes, interaction strength therefore shifts the changing balance between the two weighted contributions primarily by regulating *P*_contact_. When the attractions are sufficiently strong, patch contacts remain appreciably populated over a broader spacing range, allowing patch-contact compaction to influence the ensemble-averaged dimensions over larger spacings. The balance to the patch-noncontact contribution consequently occurs at a larger spacing, shifting the position of maximal compaction. As the attractions weaken, *P*_contact_ becomes smaller and the ensemble-averaged dimensions become increasingly governed by the patch-noncontact conformations. Under the interaction parameters considered here, *R_g,_*_noncontact_ increases with patch spacing and governs the monotonic chain expansion observed in the weak-attraction regime. How the spacing dependence of the patch-noncontact conformations changes across a broader interaction parameter space is examined in a later section.

To examine how this balance depends on patch length, we repeated the analysis using eight-residue hydrophobic patches (Fig. S5). Reducing the patch length preserved the overall progression from monotonic to nonmonotonic behavior as the interaction strengths increased, but shifted the spacing of maximal compaction toward smaller values and reduced the region of parameter space exhibiting nonmonotonic compaction. These results are consistent with larger hydrophobic patches maintaining appreciable *P*_contact_ over a wider range of patch spacings, thereby extending the spacing range over which contact-state compaction influences the ensemble.

### Effective Interaction Length Scale Regulates the Optimal Hydrophobic Spacing

Having established how interaction strength regulates the response of chain dimensions to hydrophobic patch spacing, we next asked how the spatial scale of the interactions influences this response. Whereas interaction strength controls the stability of patch contacts, the spatial extent of the interactions may determine how readily those contacts are maintained as patch spacing increases. Recent CG simulations showed that extending an effective synapsin interaction to retain its weak outer repulsive contribution altered aggregate organization.^40^ Within the Ashbaugh–Hatch potential,^41^ *σ* sets the characteristic length scale of both the excluded-volume core and the attractive interaction well. The previous simulations were performed using the CG residue-size parameter of *σ* = 0.55 nm for both patch and nonpatch residues. Increasing *σ* shifts the steric core outward and broadens the attractive well. Because these effects cannot be varied independently through *σ*, we refer to their combined spatial scale as the effective interaction length scale. To examine how this parameter regulates chain compaction, we varied it from 0.45 to 0.65 nm while keeping the remaining model parameters unchanged.

Increasing *σ* systematically shifted the spacing of maximal compaction toward larger values (Fig. 3A–C). At *σ* = 0.45 nm, the minimum occurred at a relatively short patch spacing and was observed over a limited region of the interaction-strength parameter space (Fig. 3A,D and Fig. S6). As *σ* increased, the minimum progressively shifted toward larger patch spacings and emerged across a broader range of *λ*_1_ and *λ*_2_ values (Fig. 3B,C,E,F and Fig. S7). At the largest *σ* and strongest interaction combination, *R_g_* decreased throughout the sampled spacing range, consistent with the minimum shifting beyond the largest accessible spacing. Thus, increasing the effective interaction length scale both shifts the optimal patch spacing and broadens the region of parameter space exhibiting contact-mediated compaction.

**Figure 3:**
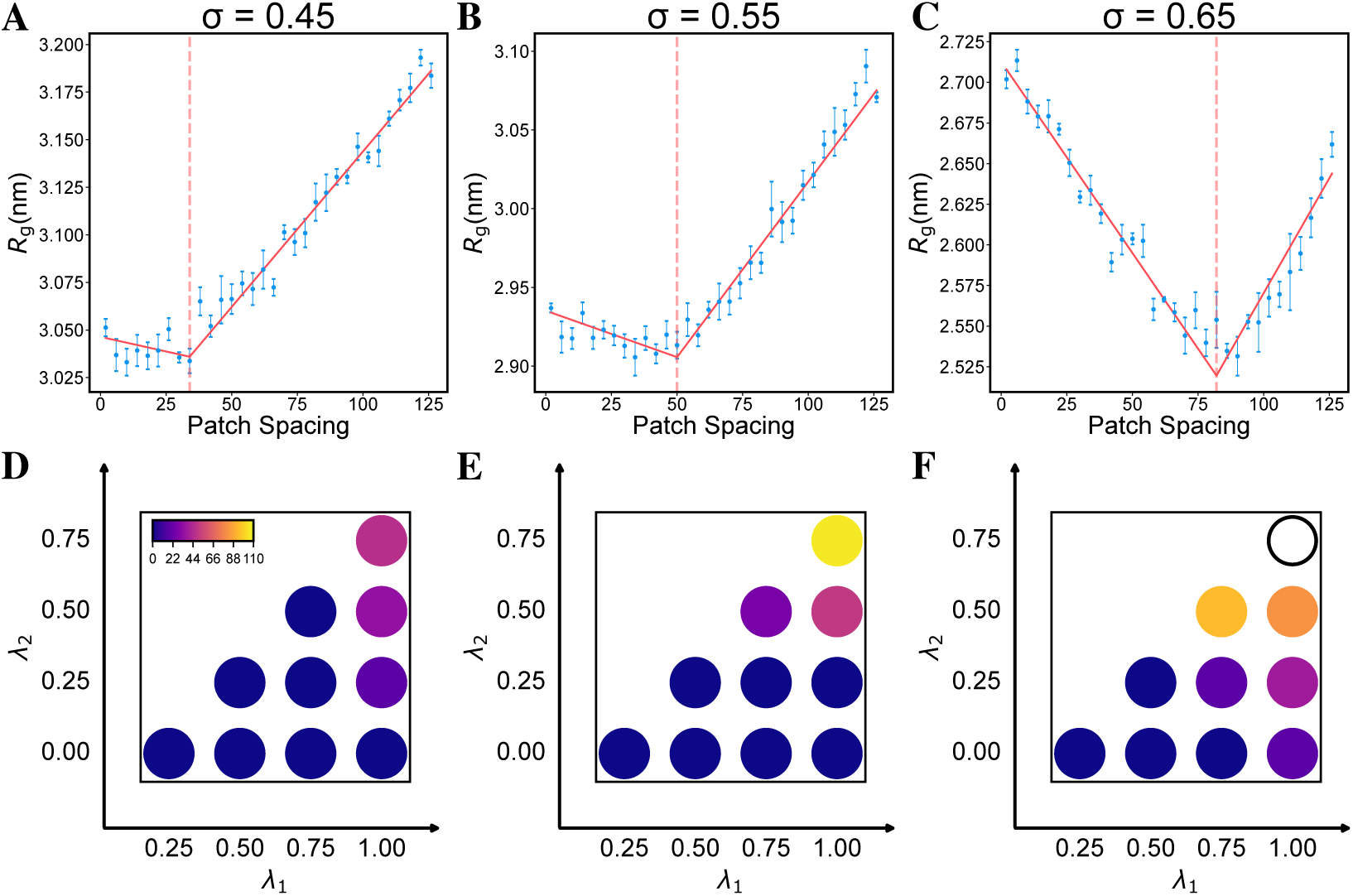
Increasing the effective interaction length scale shifts the optimal hydrophobic spacing toward larger values. (A–C) *R_g_* as a function of hydrophobic patch spacing for model peptides with a common CG residue-size parameter of *σ* = 0.45, 0.55, and 0.65 nm, respectively, at *λ*_1_ = 1.0 and *λ*_2_ = 0.5. The same *σ* was assigned to the patch and nonpatch residues. Red lines represent piecewise linear fits, and vertical dashed lines indicate the corresponding spacing of maximal compaction (see Methods). (D–F) Optimal patch spacing determined from the piecewise linear analysis across different combinations of *λ*_1_ and *λ*_2_ at *σ* = 0.45, 0.55, and 0.65 nm, respectively. Colors indicate the optimal patch spacing. Hollow circles indicate monotonic chain compaction for which no optimal spacing was assigned, whereas zero indicates monotonic chain expansion. The complete *R_g_* profiles for *σ* = 0.45 and 0.65 nm are shown in Figs. S6 and S7, respectively.

To determine the origin of this shift, we performed the same conformational-class decomposition used in Fig. 2B–D for *σ* = 0.45, 0.55, and 0.65 nm at *λ*_1_ = 1.0 and *λ*_2_ = 0.5 (Fig. S8). For all three values of *σ*, the weighted patch-contact contribution decreased with patch spacing, whereas the weighted patch-noncontact contribution increased. However, increasing *σ* shifted the change in dominance between these opposing contributions toward larger patch spacings. The class-conditioned chain dimensions retained similar qualitative spacing dependences across the three values of *σ*. In contrast, *P*_contact_ became substantially more persistent as *σ* increased. The weighted patch-contact contribution therefore remained appreciable and continued to govern the spacing response over a larger range, while the increase in the patch-noncontact contribution was delayed. These results indicate that increasing the effective interaction length scale shifts the spacing of maximal compaction primarily by maintaining patch contacts over larger sequence separations.

### Multiple Hydrophobic Patches Regulate the Optimal Spacing

The preceding sections established how interaction strength and effective interaction length scale regulate the optimal hydrophobic spacing in model peptides containing two hydrophobic patches. Naturally occurring IDPs, however, often contain multiple localized hydrophobic regions that may interact cooperatively. To determine how patch multiplicity affects sequence-dependent chain compaction, we introduced a third 12-residue hydrophobic patch at the midpoint between the original two patches while maintaining the peptide length and the remaining model parameters (Fig. 4A,B). Introducing the third patch converts a central group of nonpatch residues with interaction strength *λ*_2_ into patch residues with interaction strength *λ*_1_. The magnitude of this perturbation is therefore related to the interaction contrast, defined as Δ*λ* = *λ*_1_ − *λ*_2_. When Δ*λ* = 0, patch and nonpatch residues are physically equivalent, and the additional labeled patch does not alter the peptide.

**Figure 4:**
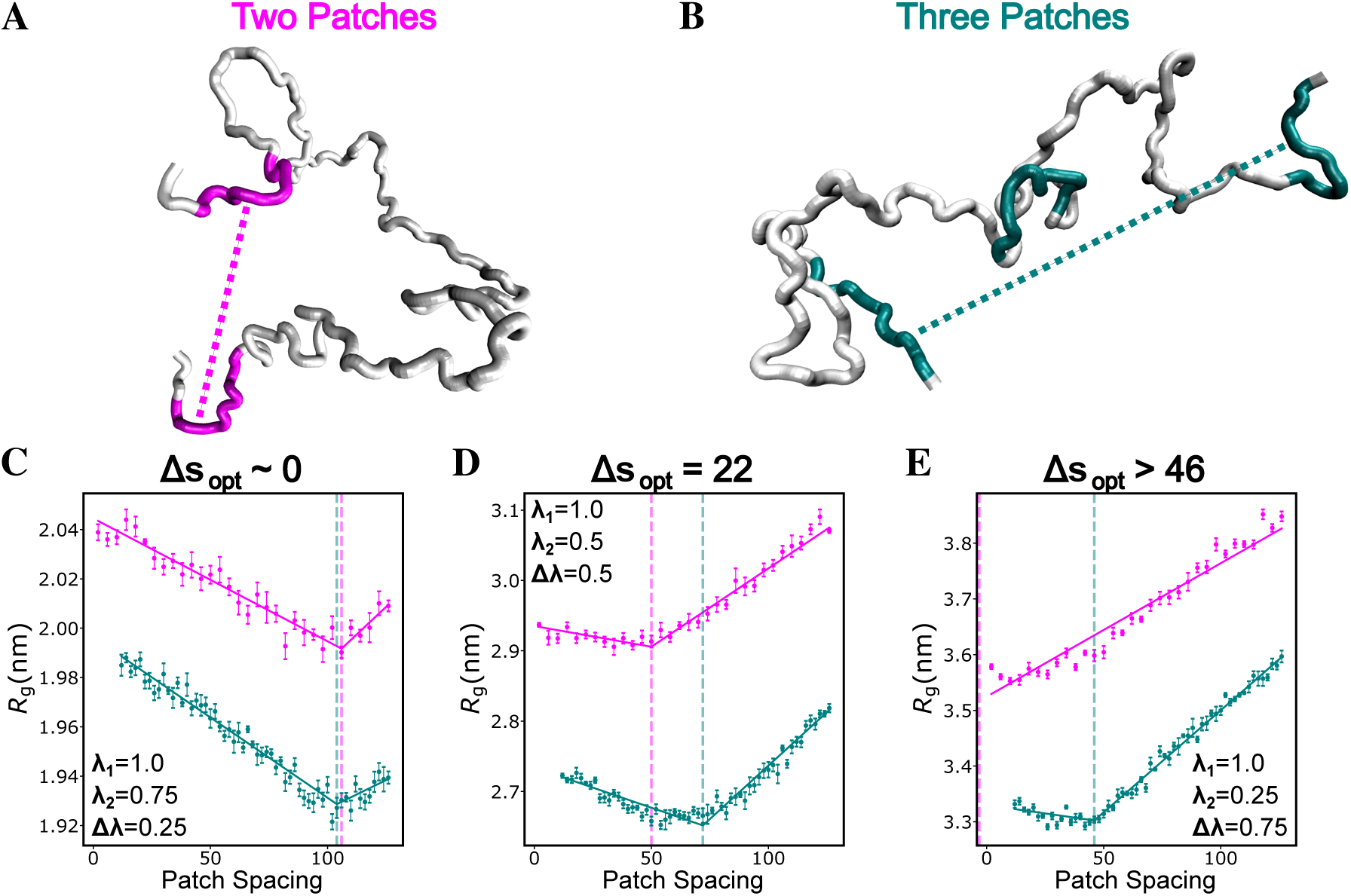
Interaction contrast controls the shift in optimal hydrophobic spacing produced by an additional patch. (A,B) Representative conformations of model peptides containing two and three 12-residue hydrophobic patches, respectively. (C–E) *R_g_* as a function of hydrophobic patch spacing for the two-patch (magenta) and three-patch (cyan) peptides at *λ*_1_ = 1.0 and *λ*_2_ = 0.75, 0.50, and 0.25, respectively. Solid lines represent piecewise linear fits, and vertical dashed lines indicate the corresponding spacing of maximal compaction (see Methods). 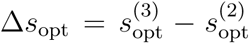 denotes the difference in optimal spacing between the three-and two-patch architectures. In panel E, Δ*s*_opt_ *>* 46 is a lower bound because 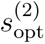 lies below the sampled range.

The effect of the third hydrophobic patch became increasingly pronounced as the interaction contrast increased (Fig. 4C–E). At fixed *λ*_1_ = 1.0, decreasing *λ*_2_ from 0.75 to 0.25 increased Δ*λ* from 0.25 to 0.75. Although the optimal patch spacing shifted toward smaller values for both architectures as *λ*_2_ decreased, this shift was substantially weaker for the three-patch peptide. Consequently, the difference between the two architectures increased from approximately zero at Δ*λ* = 0.25 to 22 residues at Δ*λ* = 0.50 and more than 46 residues at Δ*λ* = 0.75. The same relationship is apparent across the broader interaction parameter space (Fig. S9), suggesting that Δ*λ* organizes the effect of adding a third patch. The interaction contrast does not, however, determine the absolute optimal spacing by itself, because the absolute values of *λ*_1_ and *λ*_2_ also determine the overall interaction regime. These results indicate that the effect of patch multiplicity depends on how strongly the patches are differentiated from the surrounding sequence.

To determine how the additional hydrophobic patch shifts the spacing of maximal compaction, we compared the conformational-class decomposition of the two-and three-patch systems using the same contact definition for the original pair of hydrophobic patches (Fig. S10). At interaction contrasts of Δ*λ* = 0.75 and 0.50, introducing the third patch reduced the sensitivity of *P*_contact_ to patch spacing. Although the contact probability was not uniformly increased, its decay was slower, preserving contact between the original patch pair over a broader spacing range. The three-patch system also exhibited smaller class-averaged chain dimensions at these interaction contrasts, although the spacing dependences of *R_g,_*_contact_ and *R_g,_*_noncontact_ were not systematically enhanced. By contrast, at Δ*λ* = 0.25, the contact probabilities, class-averaged dimensions, and weighted contributions of the two architectures largely overlapped, indicating that the effect of the additional patch diminishes as the interaction contrast decreases. Thus, at larger interaction contrasts, the smaller class-averaged dimensions contribute to the overall compaction of the three-patch system, whereas the shift in maximal compaction arises primarily from the slower redistribution of conformational population from the patch-contact to the more expanded patch-noncontact class. By sustaining the influence of patch-contact conformations over a broader spacing range, the additional patch shifts maximal compaction toward larger patch spacings.

### A Unified Framework Connects Three Distinct Spacing Responses

The preceding sections examined how interaction strength, effective interaction length scale, and patch architecture regulate chain dimensions within predominantly attractive interaction regimes. We next asked whether the same conformational-class framework could describe conditions in which steric excluded volume interactions dominate. To access both regimes, we assigned different CG residue-size parameters to the patch and nonpatch residues, denoted *σ*_1_ and *σ*_2_, respectively, while systematically varying *λ*_1_ and *λ*_2_. When *σ*_1_ *> σ*_2_, the patch residues introduce an excluded-volume contrast at weak attraction but become increasingly attractive relative to the nonpatch residues as *λ*_1_ increases. This combined parameter scan therefore connects steric-dominated and attraction-dominated sequence patterning within the same model.

For the representative combination *σ*_1_ = 0.65 and *σ*_2_ = 0.45 nm, increasing the patch attraction produced a systematic progression among three spacing responses (Fig. 5). At weak patch attractions (*λ*_1_ = 0, 0.25, and 0.5), *R_g_* decreased monotonically with patch spacing. For off-diagonal conditions with *λ*_1_ *> λ*_2_, increasing *λ*_1_ to 0.75 produced monotonic chain expansion, whereas further increasing *λ*_1_ to 1.0 produced a nonmonotonic response. In contrast, conditions along the *λ*_1_ = *λ*_2_ diagonal remained monotonically compacting across the sampled interaction strengths. The same three responses were observed for additional combinations of *σ*_1_ and *σ*_2_ (Figs. S11 and S12), although the boundaries between the regimes shifted with the residue-size parameters.

**Figure 5:**
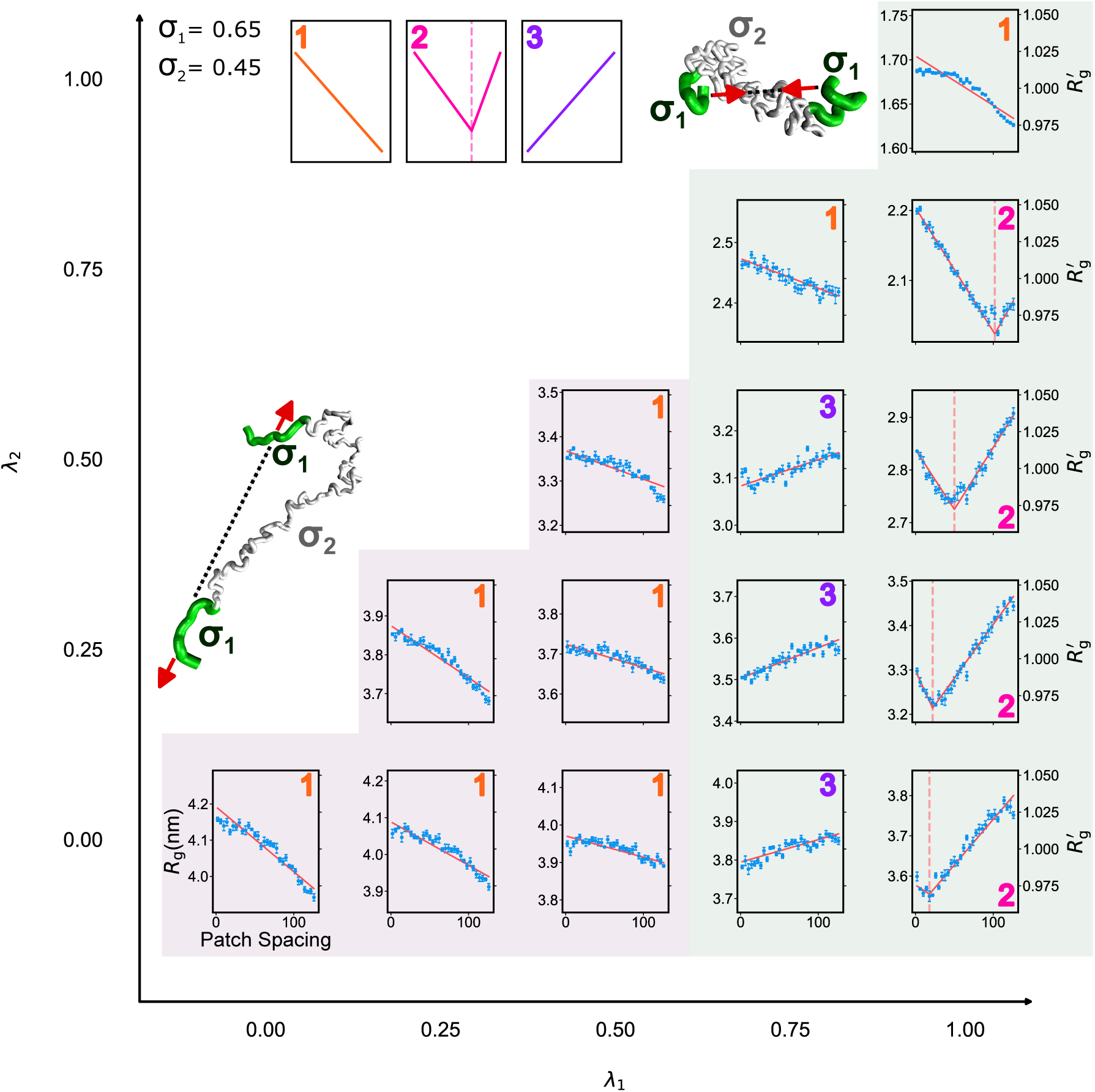
A unified framework connects three distinct spacing responses. *R_g_* as a function of hydrophobic patch spacing for model peptides with patch and nonpatch residue-size parameters of *σ*_1_ = 0.65 and *σ*_2_ = 0.45 nm, respectively, across the sampled combinations of patch (*λ*_1_) and nonpatch (*λ*_2_) interaction strengths. The numbered labels identify monotonic chain compaction (1, orange), nonmonotonic compaction (2, magenta), and monotonic chain expansion (3, purple). Red lines represent fits used to classify the spacing responses, and vertical dashed lines indicate the spacing of maximal compaction for nonmonotonic profiles. The right axes show *R^′^* = *R_g_/R_g_*, where *R_g_* is the mean *R_g_* across all sampled patch spacings for the corresponding combination of interaction parameters. Representative conformations and schematics illustrate the dominant steric and attractive interactions. Light-purple and light-green shading indicate steric-dominated and attraction-dominated regions, respectively.

The conformational-class decomposition established in Fig. 2 provides a direct explanation for these responses (Figs. S13–S15). At weak patch attractions, *P*_contact_ decreased rapidly with patch spacing, and the ensemble became dominated by patch-noncontact conformations over most of the spacing range (Figs. S13 and S14). In this steric-dominated regime, *R_g,_*_noncontact_ decreased with patch spacing (Fig. S15), consistent with progressive relief of the steric frustration produced by clustering the larger patch residues. The resulting decrease in the patch-noncontact contribution produced monotonic chain compaction.

At intermediate patch attractions, the spacing dependence of *R_g,_*_noncontact_ changed qualitatively. Rather than decreasing with patch spacing as in the weak-attraction regime, *R_g,_*_noncontact_ became approximately constant or weakly increasing (Fig. S15). The patch-noncontact conformations therefore no longer produced the sterically driven compaction observed at weaker attractions. Because *P*_noncontact_ continued to increase with spacing, the weighted patch-noncontact contribution increased and dominated the ensemble response, producing monotonic chain expansion. This behavior recovers the monotonic expansion regime observed within the attractive interaction parameter space of Fig. 2.

At strong patch attraction, *P*_contact_ remained appreciable over a broader spacing range, and the decreasing patch-contact contribution continued to influence the ensemble response. At smaller spacings, the decrease in the patch-contact contribution dominated and caused the chain to compact. At larger spacings, the increasing patch-noncontact contribution became dominant and caused the chain to expand, producing the nonmonotonic response. This reproduces the contact-mediated mechanism established in Fig. 2, despite the difference between the patch and nonpatch residue-size parameters. Thus, increasing the patch attraction shifts the system from a steric-dominated compaction regime into the monotonic and nonmonotonic attractive regimes established in Fig. 2.

These results generalize the conformational-class decomposition established in Fig. 2 into a unified framework spanning steric-and attraction-dominated regimes. Within this framework, the interaction parameters regulate the relative magnitudes and spacing dependences of the weighted patch-contact and patch-noncontact contributions. A monotonic decrease in *R_g_* can arise through two distinct microscopic mechanisms: progressive compaction of the dominant patch-noncontact conformations in the steric-dominated regime, or sustained dominance of the decreasing patch-contact contribution under strongly attractive conditions. The same macroscopic spacing response can therefore reflect distinct conformational origins that can be resolved through analysis of the weighted contributions.

## Conclusion

Hydrophobic patch spacing is not a uniformly monotonic determinant of chain dimensions of intrinsically disordered proteins. At fixed amino acid composition, FUS-derived sequence variants exhibited maximal compaction at an intermediate spacing between hydrophobic patches. Analysis using simplified model peptides showed that this nonmonotonic response emerges only within sufficiently attractive interaction regimes. The spacing of maximal compaction shifted toward larger values with increasing attraction strength, effective interaction length scale, and patch length. Introducing an additional hydrophobic patch produced a further architecture-dependent shift whose magnitude increased with the interaction contrast between the patch and nonpatch residues. These results establish that hydrophobic sequence organization depends not only on pairwise sequence separation but also on the interaction regime.

The conformational-class decomposition unifies the observed spacing responses through the weighted patch-contact and patch-noncontact contributions. Within attractive regimes, the patch-contact contribution decreases with spacing, whereas the patch-noncontact contribution increases, and their relative slopes determine whether the chain expands, compacts, or exhibits a nonmonotonic response. When steric interactions dominate, *R_g,_*_noncontact_ instead decreases as separating the larger patch residues relieves steric frustration. A monotonic decrease in *R_g_* can therefore arise either from compaction of the dominant patch-noncontact conformations or from sustained dominance of the decreasing patch-contact contribution.

Although electrostatic interactions were not modeled explicitly, the steric-dominated and strongly attractive limits provide qualitative analogues of repulsive and attractive charge interactions, respectively. Sequence Charge Decoration encodes both attractive and repulsive electrostatic interactions using a common spacing dependence. ^16^ Our results suggest that similar macroscopic spacing responses may nevertheless arise from distinct microscopic mechanisms. For hydrophobic interactions, Sequence Hydropathy Decoration captures the monotonic limit of hydrophobic interactions,^27^ whereas the nonmonotonic regime identified here lies beyond its fixed monotonic dependence on spacing. The framework predicts that weakening hydrophobic attraction should eliminate the spacing-dependent maximal compaction, whereas stronger attraction, larger patches, or additional high-contrast patches should shift maximal compaction toward larger spacings.

## Methods

### Coarse-grained molecular dynamics simulations

Single-chain coarse-grained (CG) molecular dynamics simulations were performed using the CALVADOS2 model^33^ implemented in OpenMM.^42^ All systems were simulated at 310 K using a Langevin thermostat with a friction coefficient of 0.01 ps*^−^*^1^ and a time step of 10 fs. Each system was simulated for 2.0 µs, with the first 0.5 µs discarded as equilibration and the remaining 1.5 µs used for analysis. The FUS-derived sequence variants were simulated using the default CALVADOS2 residue parameters. For the simplified model peptides, the patch and nonpatch residues were assigned interaction parameters *λ*_1_ and *λ*_2_, respectively, and, where indicated, residue-size parameters *σ*_1_ and *σ*_2_. When a single *σ* value is reported, the same value was assigned to all residues. All other force-field terms and functional forms were retained from the original model.

Patch spacing, *s*, was defined as the number of residues between the nearest edges of the two reference hydrophobic patches. To vary *s* while preserving the chain length and composition, the two patches were displaced simultaneously and symmetrically in opposite directions along the sequence. For the 163-residue FUS-derived sequences, patch spacing was sampled at *s* = 1, 3, 5, 7*, …*, corresponding to two-residue increments. For the 150-residue model peptides, patch spacing was sampled at *s* = 2, 6, 10, 14*, …*, corresponding to four-residue increments. For the three-patch peptides, *s* continued to denote the spacing between the two original reference patches, while the additional patch was positioned at their midpoint.

Reported *R_g_* values were calculated from the complete production trajectory. To estimate statistical uncertainty, each production trajectory was divided into five equal-length blocks of 0.3 µs. Error bars represent the standard error of the mean calculated from the five block-averaged *R_g_* values.

### Piecewise linear analysis

To identify the hydrophobic patch spacing corresponding to maximal chain compaction, each *R_g_* profile was fit using both a single-line model and a continuous piecewise linear model. The piecewise model was defined as

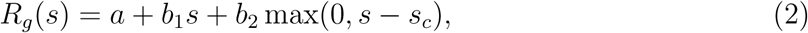

where *s* is the patch spacing and *s_c_* is the breakpoint. The slopes below and above the breakpoint are *b*_1_ and *b*_1_ + *b*_2_, respectively. Fits were performed by weighted least squares, with each data point weighted by the inverse square of its standard error. The breakpoint *s_c_* was treated as a continuous fitting parameter constrained to the sampled spacing range, with at least three data points required on each side. The single-line and piecewise models were compared using the Akaike information criterion (AIC),^43^ with two fitted parameters assigned to the single-line model and four to the piecewise model, including the breakpoint. A profile was classified as nonmonotonic when the piecewise model had a lower AIC than the single-line model and exhibited a negative slope below the breakpoint and a positive slope above it. For profiles satisfying these criteria, the breakpoint was defined as the optimal hydrophobic spacing, *s*_opt_. Profiles not satisfying these criteria were classified as monotonic, and no numerical *s*_opt_ was assigned. A monotonically expanding profile indicates that *s*_opt_ lies below the sampled spacing range, whereas a monotonically compacting profile indicates that *s*_opt_ lies above the sampled range.

### Conformational-class decomposition

Conformations were classified according to the minimum distance between CG residues belonging to the two reference hydrophobic patches. For the three-patch peptides, the same pair of patches used in the corresponding two-patch system was used for classification, and the additional central patch was not included in the contact criterion. A conformation was assigned to the patch-contact class if any residue pair spanning the two reference patches was separated by less than 1.0 nm and to the patch-noncontact class otherwise. For each class, the population fraction and class-averaged *R_g_* were calculated from the corresponding trajectory frames. Statistical uncertainties were estimated by dividing each production trajectory into five equal, nonoverlapping temporal blocks and recalculating the population fraction and class-averaged *R_g_* within each block. Error bars represent the standard error of the mean across the five block estimates. Conformational classes containing fewer than 50 frames were excluded because of insufficient sampling.

## Supporting Information Available

Supporting figures illustrating additional parameters.

## Acknowledgement

This work is supported by the National Institutes of Health (R35GM146814, W.Z.). The authors also acknowledge support from the ASU SCENE program (H.S.) and from Research Computing at Arizona State University.

