## Supplementary figures. for "Hydrophobic Patch Spacing Produces Nonmonotonic Compaction in Intrinsically Disordered Proteins"

### Supporting Information for: "Hydrophobic Patch Spacing Produces Nonmonotonic Compaction in Intrinsically Disordered Proteins"

Henry Silvernail[1], Wangfei Yang[2], and Wenwei Zheng[2,3a]

[1] Brophy College Preparatory, Phoenix, AZ 85012, USA

[2] College of Integrative Sciences and Arts, Arizona State University, Mesa, AZ 85212, USA

[3] Center for Biological Physics, Arizona State University, Tempe, AZ 85282, USA

[a]Electronic mail:

### 1 Supplementary Figures

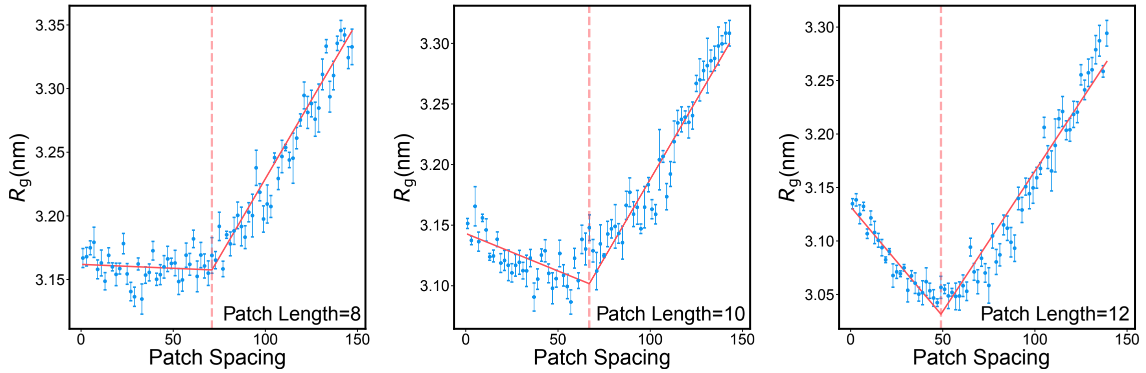


Figure S1: **Hydrophobic patch length modulates the nonmonotonic dependence of chain compaction on patch spacing.** Radius of gyration ($R_g$) as a function of hydrophobic patch spacing for composition-preserving FUS-derived sequence variants containing two tyrosine patches of 8, 10, or 12 residues. For the 8- and 10-residue patch series, 16 and 20 tyrosines, respectively, were randomly selected to form the two patches. The remaining tyrosines were retained at their original sequence positions, and the same selection was used for all patch spacings within each series. The red lines show piecewise linear fits, and the vertical dashed lines indicate the fitted patch spacings of maximal compaction.

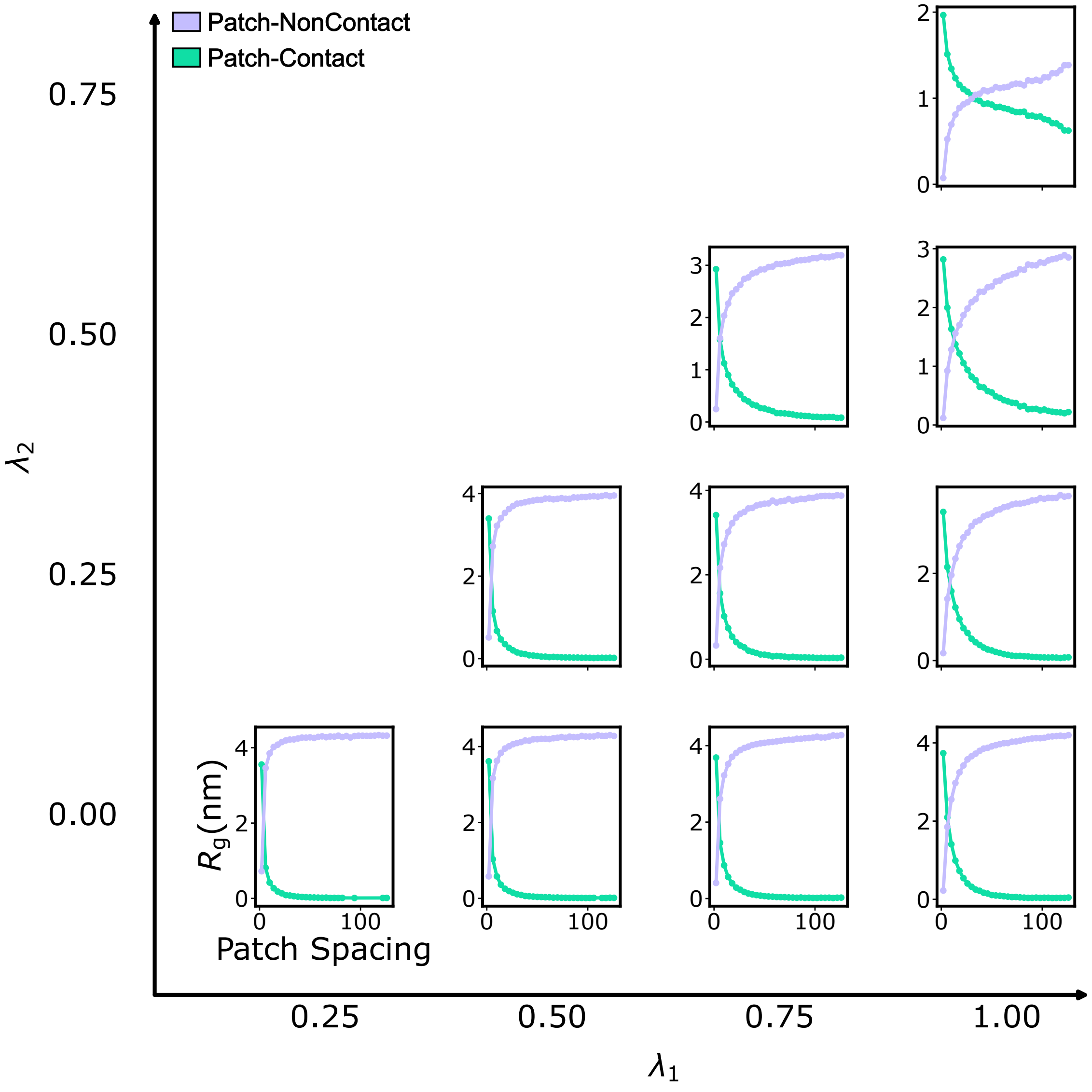


Figure S2: **Weighted contributions of the patch-contact and patch-noncontact conformational classes to the ensemble-averaged $R_g$.** Weighted contributions of the patch-contact (turquoise) and patch-noncontact (light purple) conformational classes, calculated as $P_{\text{contact}}R_{g,\text{contact}}$ and $P_{\text{noncontact}}R_{g,\text{noncontact}}$, respectively, as a function of hydrophobic patch spacing. The sum of the two contributions gives the ensemble-averaged $R_g$ in Fig. 2.

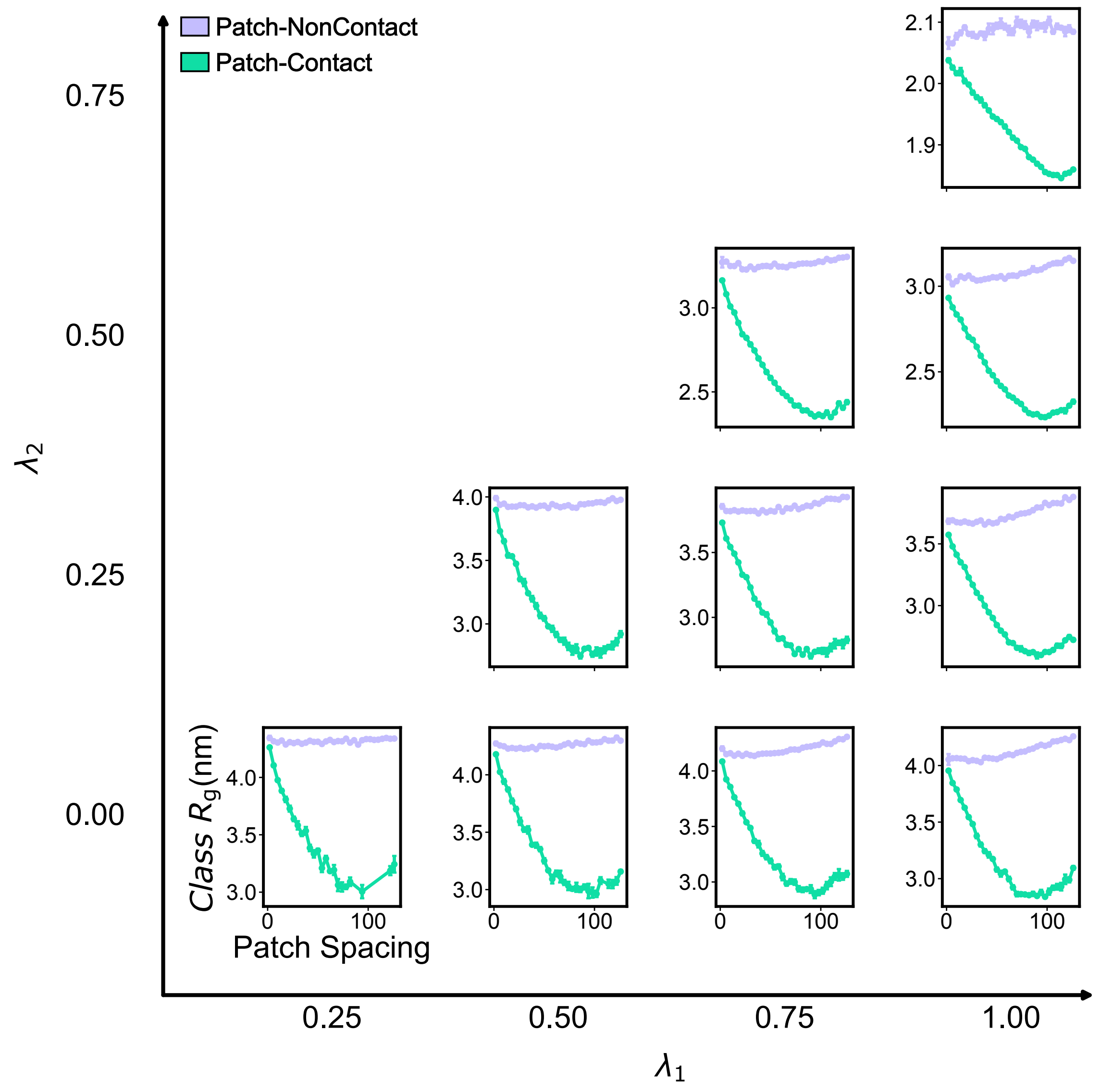


Figure S3: **Class-averaged $R_g$ of the patch-contact and patch-noncontact conformational classes.** Average $R_g$ of the patch-contact (turquoise) and patch-noncontact (light purple) conformational classes as a function of hydrophobic patch spacing.

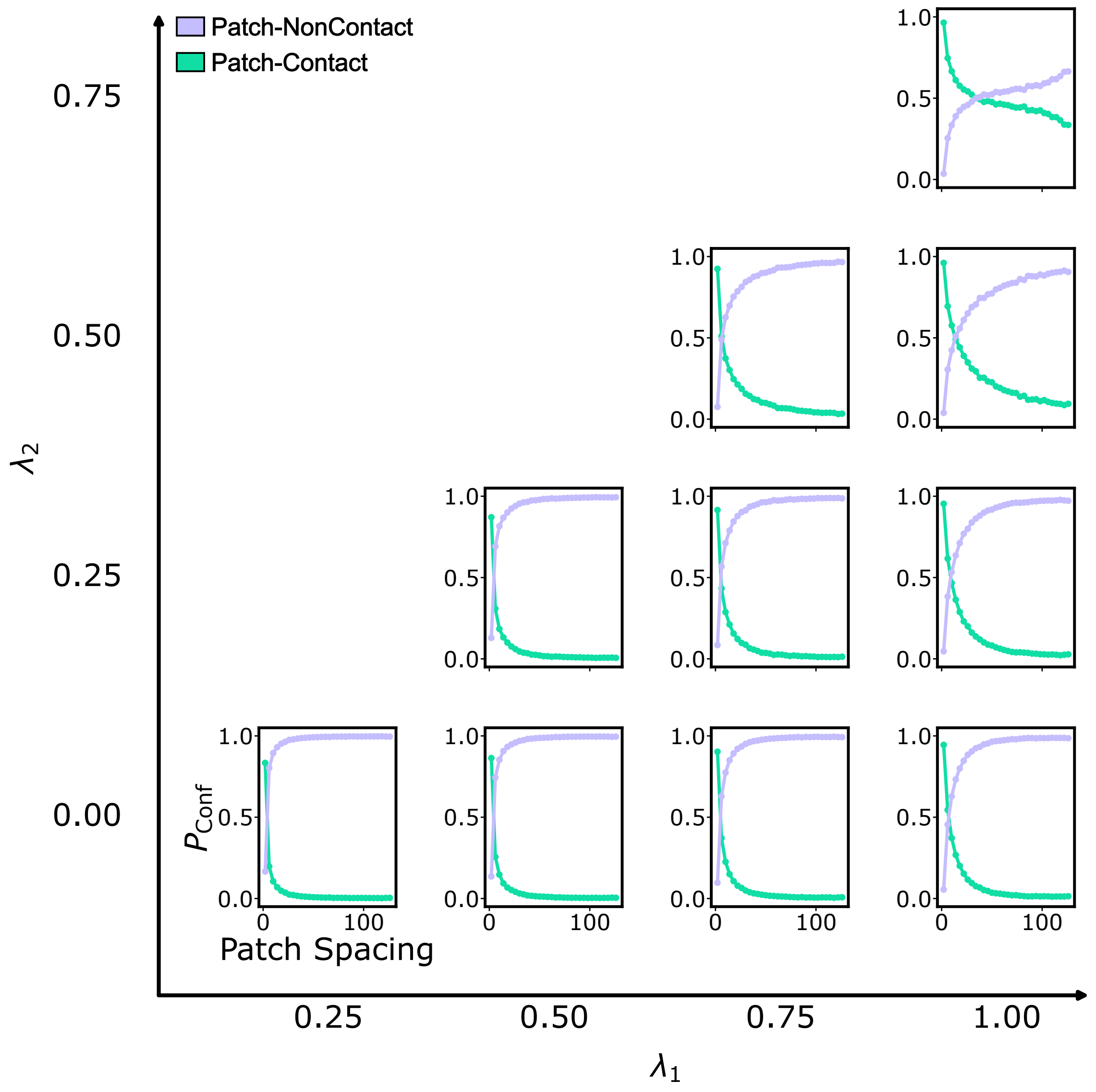


Figure S4: **Populations of the patch-contact and patch-noncontact conformational classes.** Populations of the patch-contact (turquoise) and patch-noncontact (light purple) conformational classes as a function of hydrophobic patch spacing.

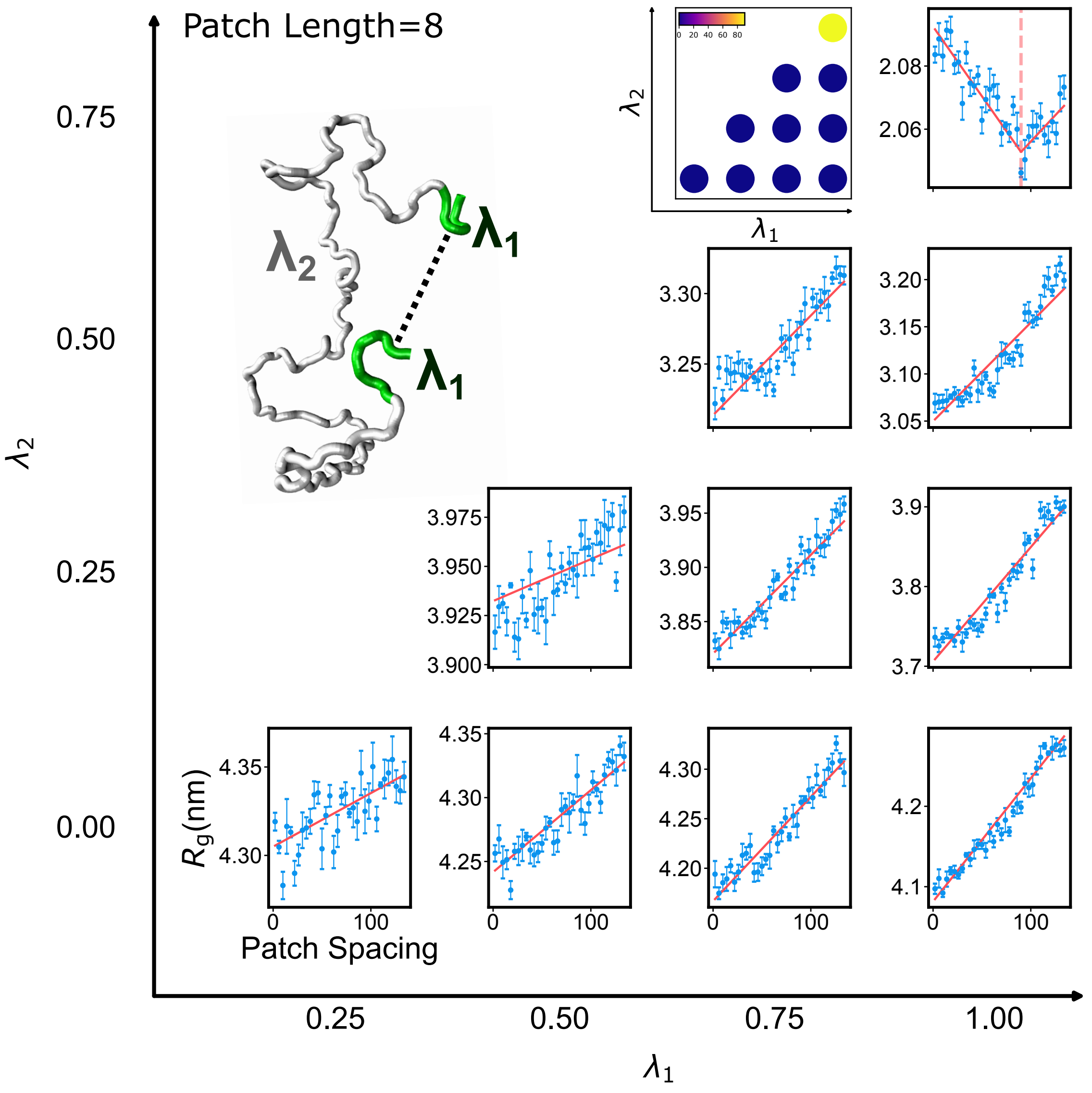


Figure S5: **Interaction strength determines the emergence of nonmonotonic compaction for eight-residue hydrophobic patches.** $R_g$ as a function of hydrophobic patch spacing for 150-residue model peptides containing two eight-residue hydrophobic patches, shown for different combinations of patch ($\lambda_1$) and nonpatch ($\lambda_2$) interaction strengths. Red lines represent piecewise linear fits used to determine the spacing of maximal compaction (see Methods). The parameter-space inset summarizes the optimal patch spacing determined from the piecewise linear analysis, with zero indicating monotonic chain expansion for which no optimal spacing was assigned. A representative conformation is shown with the hydrophobic patches in green and the nonpatch residues in gray.

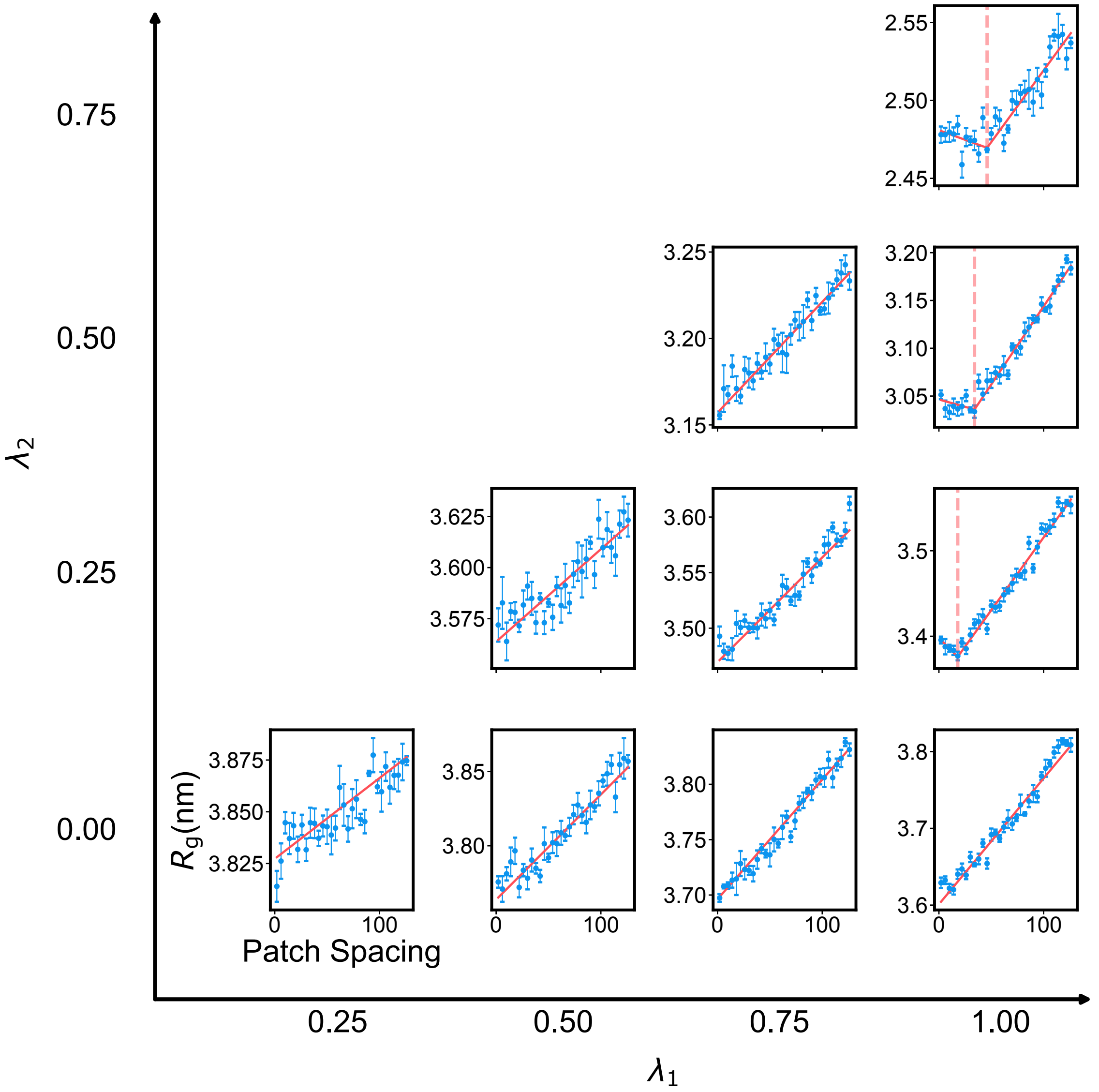


Figure S6: $R_g$ **profiles at** $\sigma = 0.45$ **nm.** $R_g$ as a function of hydrophobic patch spacing for model peptides with different combinations of patch ($\lambda_1$) and nonpatch ($\lambda_2$) interaction strengths. A common $\sigma = 0.45$ nm was assigned to all residues. Red lines represent piecewise linear fits used to determine the spacing of maximal compaction, and vertical dashed lines indicate the fitted minima (see Methods). The corresponding parameter-space summary is shown in Fig. 3D.

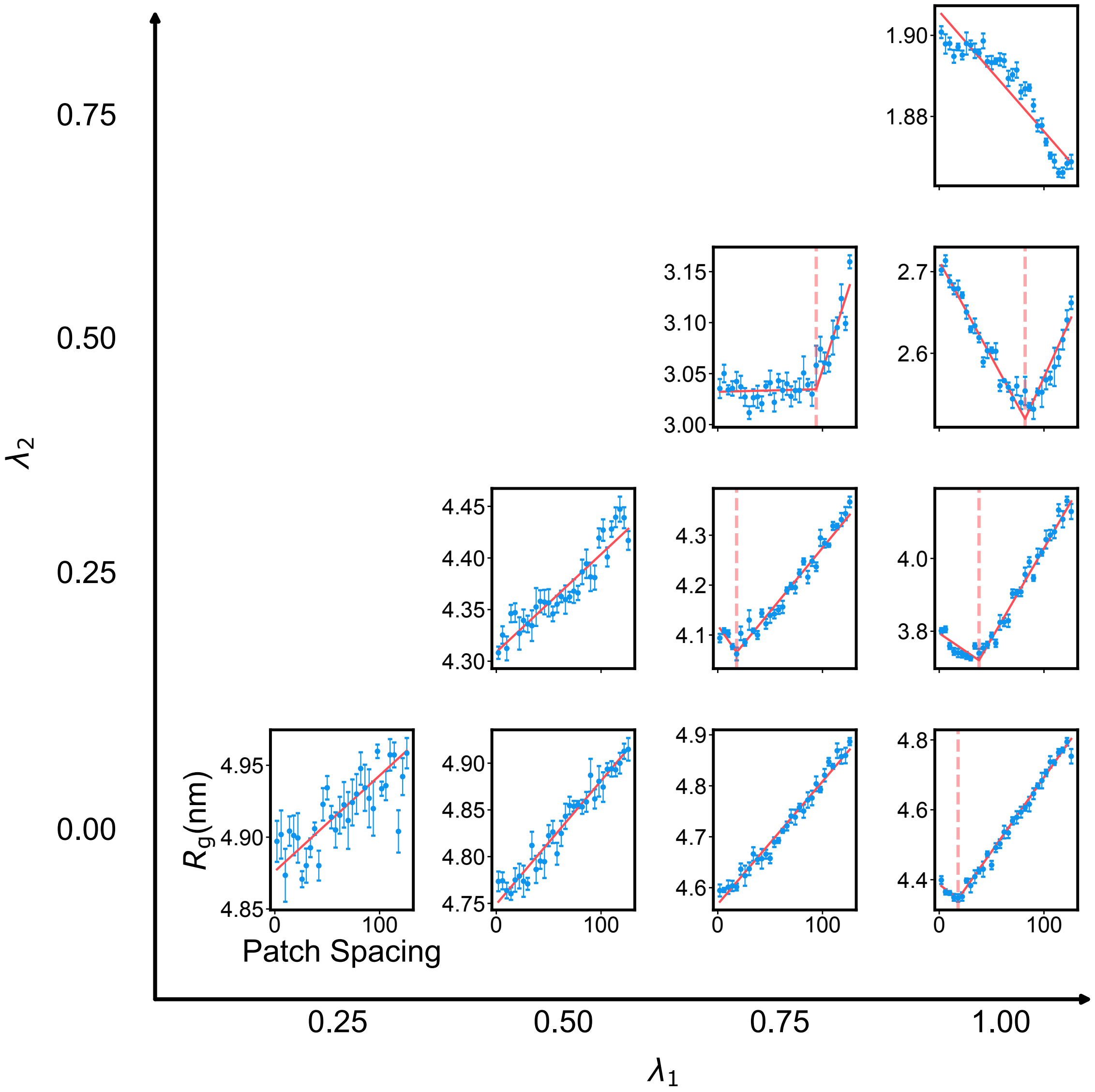


Figure S7: $R_g$ **profiles at** $\sigma = 0.65$ **nm.** $R_g$ as a function of hydrophobic patch spacing for model peptides with different combinations of patch ($\lambda_1$) and nonpatch ($\lambda_2$) interaction strengths. A common $\sigma = 0.65$ nm was assigned to all residues. Red lines represent piecewise linear fits used to determine the spacing of maximal compaction, and vertical dashed lines indicate the fitted minima (see Methods). The corresponding parameter-space summary is shown in Fig. 3F.

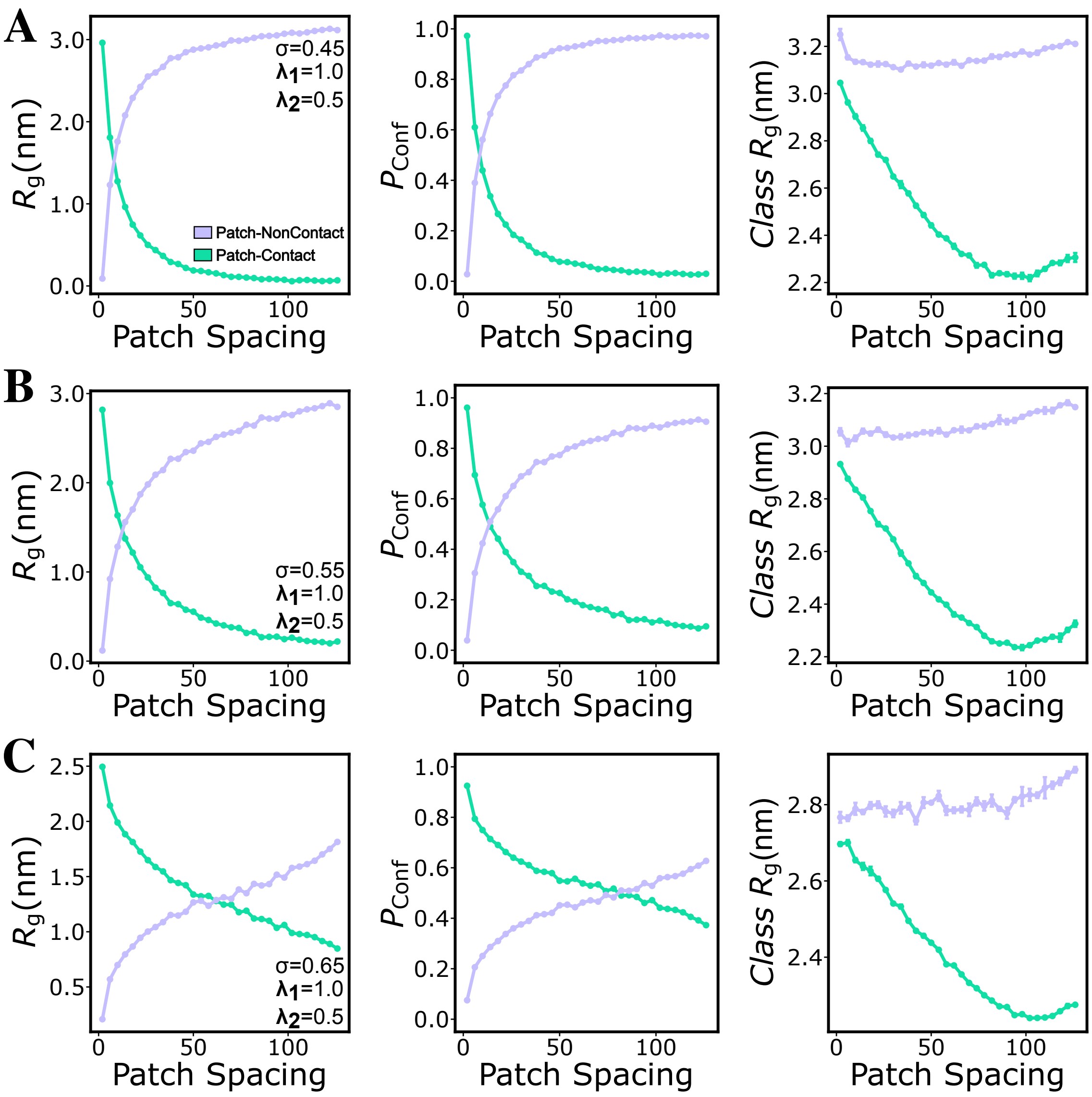


Figure S8: **Effect of the effective interaction length scale on the conformational-class contributions.** (A–C) Conformational-class decomposition at $\sigma = 0.45$, 0.55, and 0.65 nm, respectively, for $\lambda_1 = 1.0$ and $\lambda_2 = 0.5$. The left column shows the weighted contributions of the patch-contact (turquoise) and patch-noncontact (light purple) classes to the ensemble-averaged $R_g$. The middle column shows their population fractions, and the right column shows their class-averaged $R_g$.

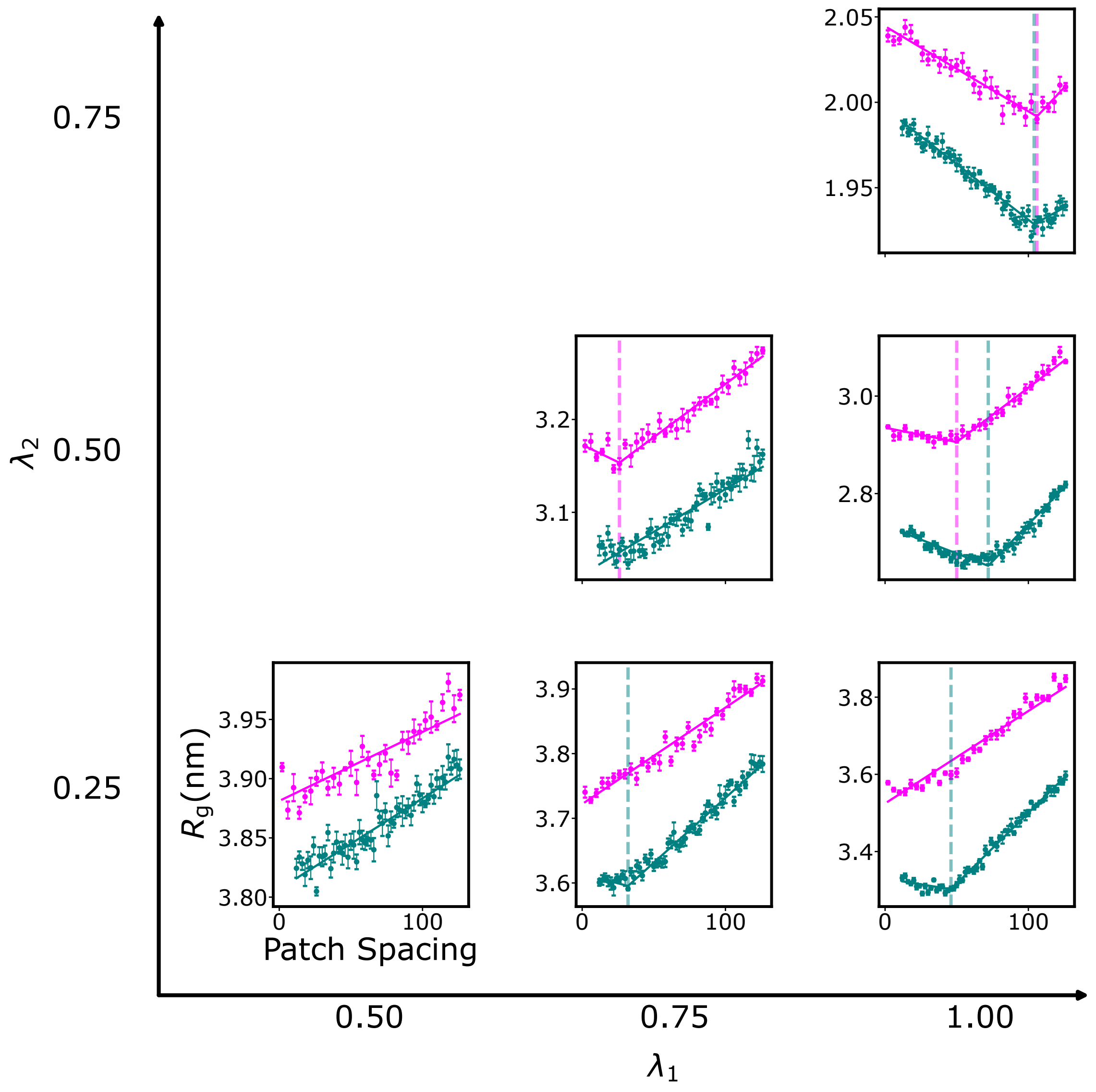


Figure S9: **Comparison of two- and three-patch model peptides across the interaction parameter space.** $R_g$ as a function of hydrophobic patch spacing for model peptides containing two (magenta) or three (cyan) 12-residue hydrophobic patches. Columns and rows correspond to increasing patch ($\lambda_1$) and nonpatch ($\lambda_2$) interaction strengths, respectively. Solid lines represent piecewise linear fits, and vertical dashed lines indicate the corresponding spacing of maximal compaction (see Methods).

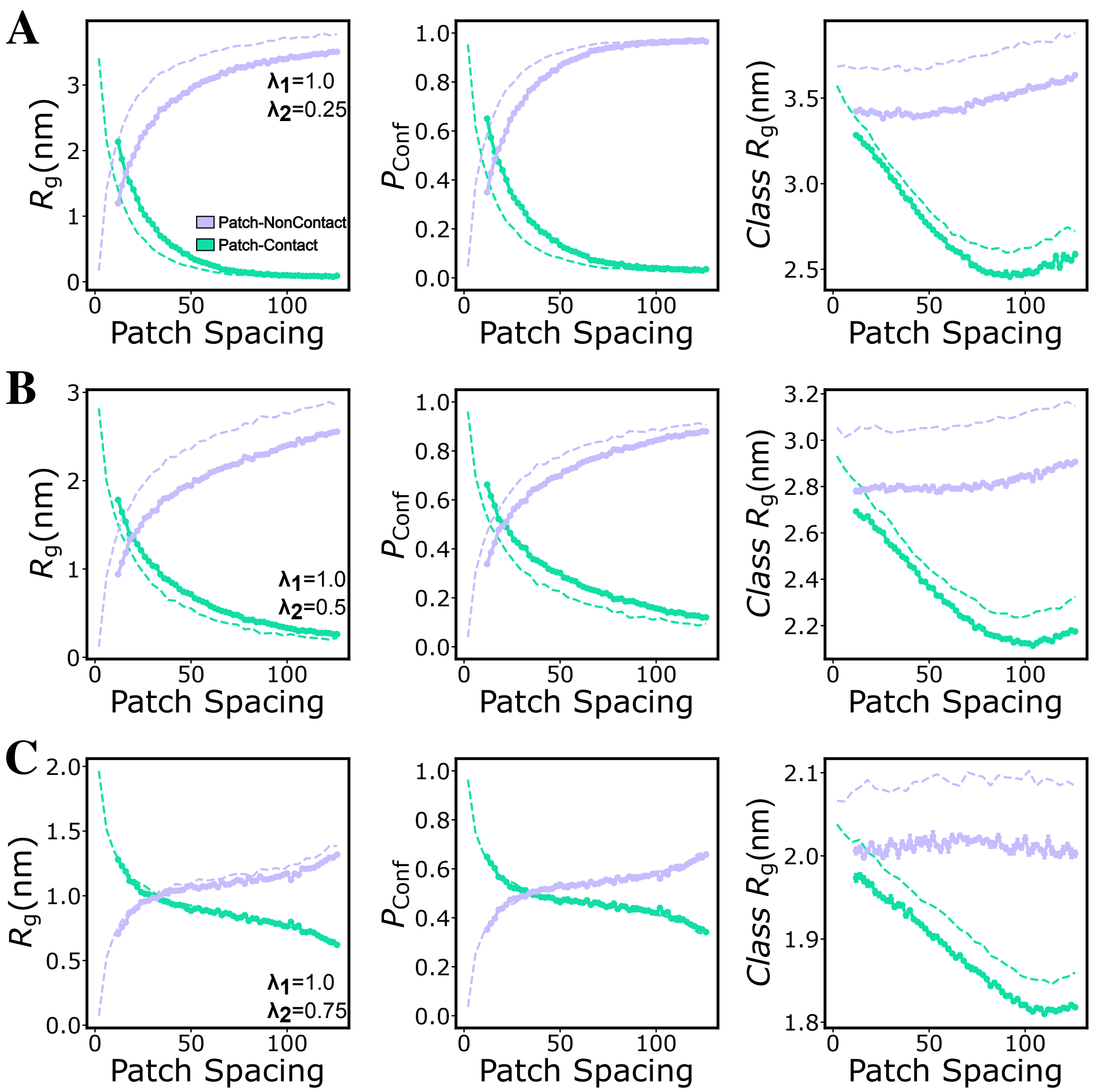


Figure S10: **Conformational-class comparison of two- and three-patch model peptides.** (A–C) Conformational-class decompositions at $\lambda_1 = 1.0$ and $\lambda_2 = 0.25$, 0.50, and 0.75, respectively. The left column shows the weighted $R_g$ contributions of the patch-contact (turquoise) and patch-noncontact (light purple) classes. The middle column shows the population fractions of the two conformational classes, and the right column shows their class-averaged $R_g$. Solid and dashed lines represent the three- and two-patch peptides, respectively.

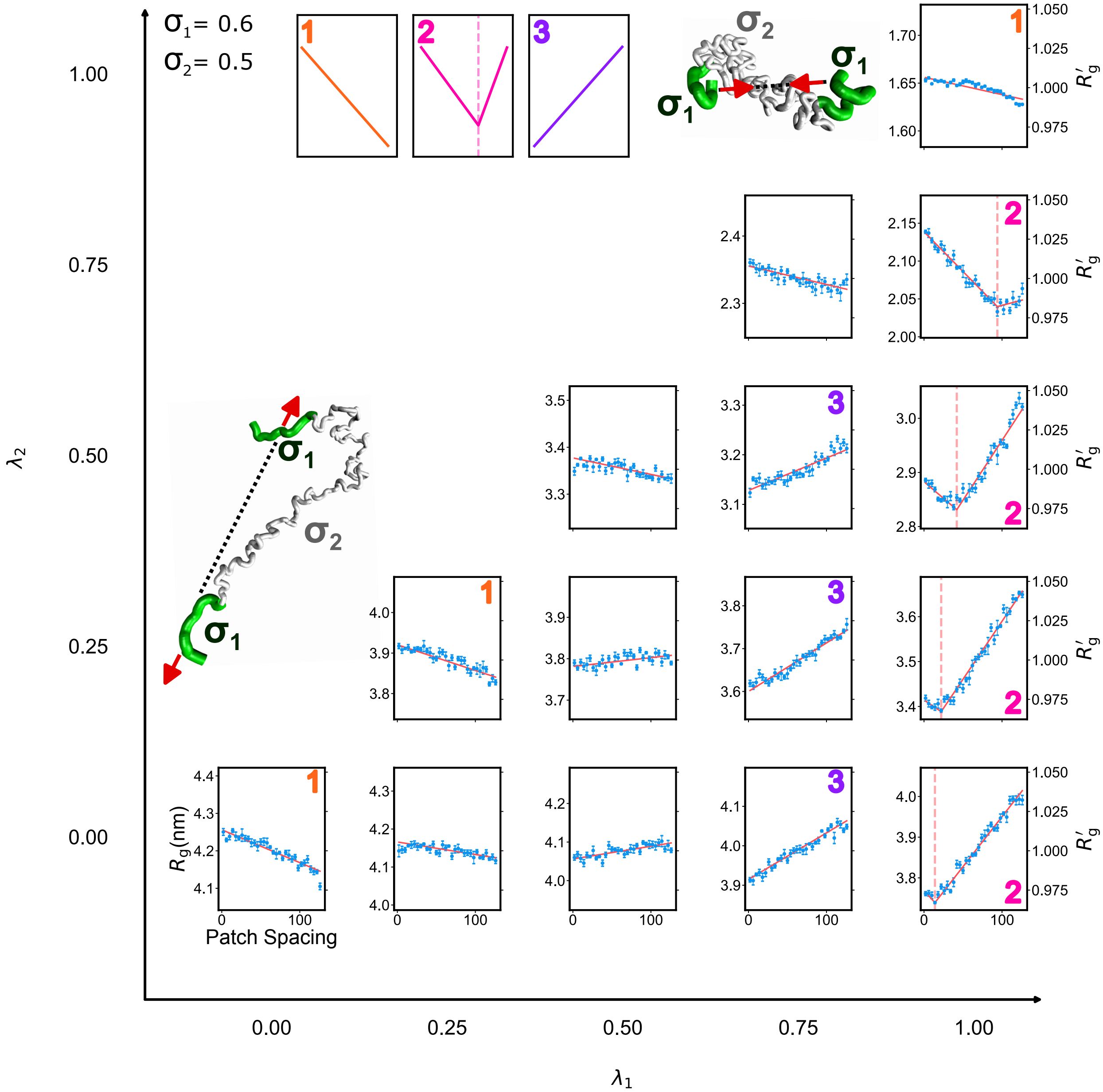


Figure S11: **Three spacing responses for $\sigma_1 = 0.60$ and $\sigma_2 = 0.50$ nm.** $R_g$ as a function of hydrophobic patch spacing across the sampled combinations of patch ($\lambda_1$) and nonpatch ($\lambda_2$) interaction strengths. The numbered labels identify monotonic chain compaction (1, orange), nonmonotonic compaction (2, magenta), and monotonic chain expansion (3, purple). Red lines represent fits used to classify the spacing responses, and vertical dashed lines indicate the spacing of maximal compaction for nonmonotonic profiles. The right axes show $R_g' = R_g/\overline{R_g}$, where $\overline{R_g}$ is the mean $R_g$ across all sampled patch spacings for the corresponding combination of interaction parameters.

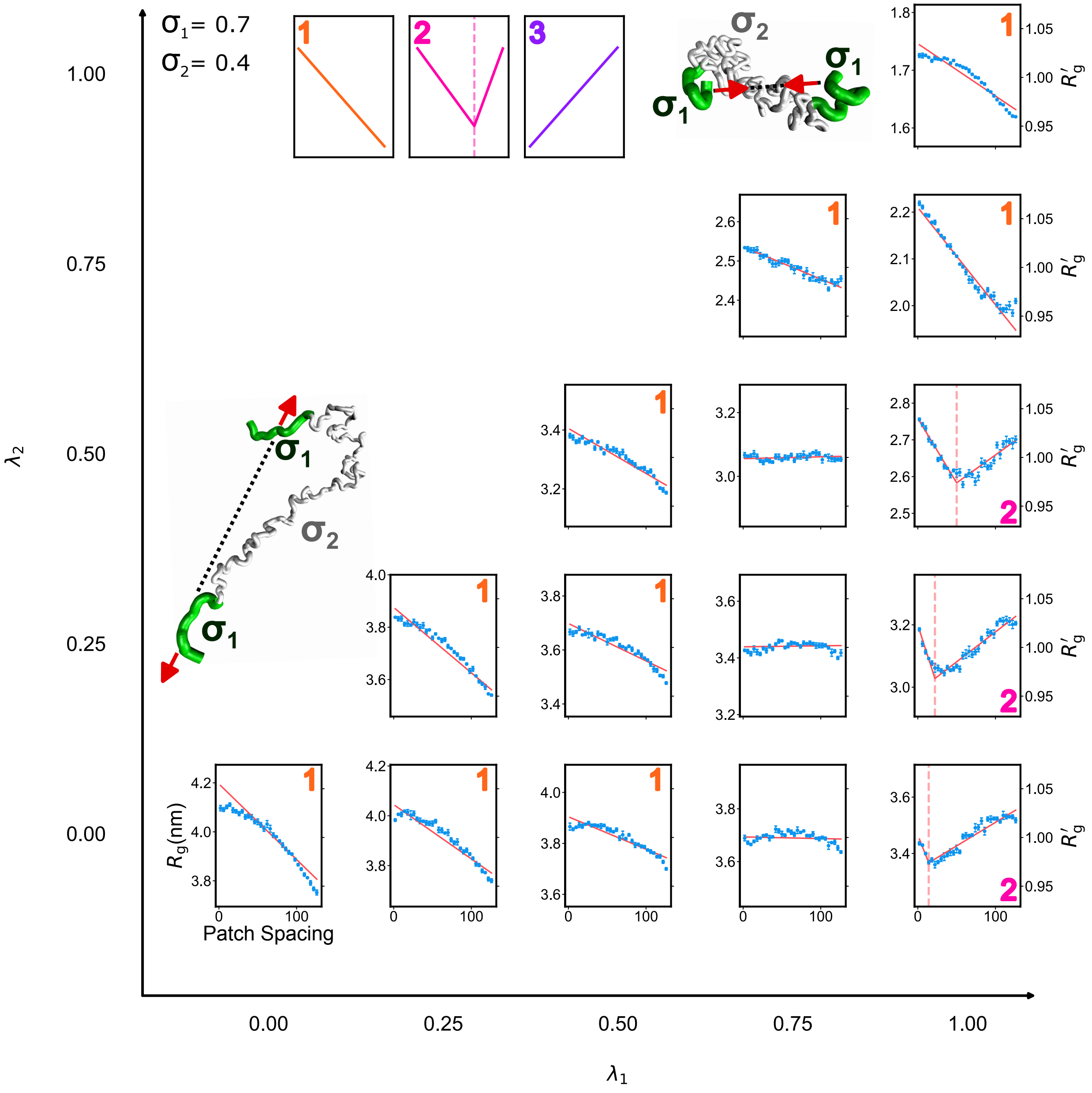


Figure S12: **Three spacing responses for $\sigma_1 = 0.70$ and $\sigma_2 = 0.40$ nm.** $R_g$ as a function of hydrophobic patch spacing across the sampled combinations of patch ($\lambda_1$) and nonpatch ($\lambda_2$) interaction strengths. The numbered labels identify monotonic chain compaction (1, orange), nonmonotonic compaction (2, magenta), and monotonic chain expansion (3, purple). Red lines represent fits used to classify the spacing responses, and vertical dashed lines indicate the spacing of maximal compaction for nonmonotonic profiles. The right axes show $R_g' = R_g/\overline{R_g}$, where $\overline{R_g}$ is the mean $R_g$ across all sampled patch spacings for the corresponding combination of interaction parameters.

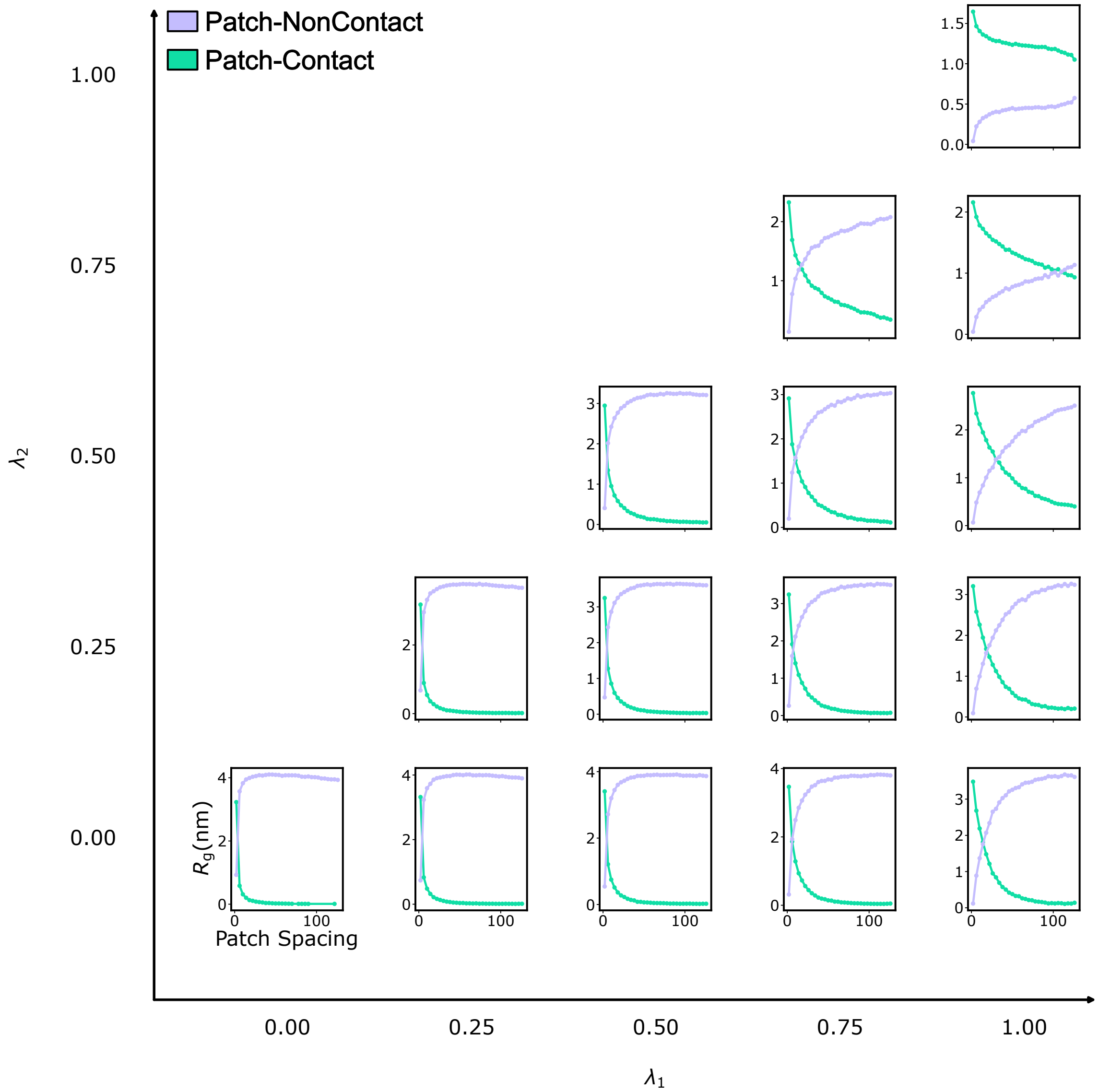


Figure S13: **Weighted conformational-class contributions for $\sigma_1 = 0.65$ and $\sigma_2 = 0.45$ nm.** Weighted $R_g$ contributions of the patch-contact (turquoise) and patch-noncontact (light purple) conformational classes as a function of hydrophobic patch spacing across the sampled combinations of patch ($\lambda_1$) and nonpatch ($\lambda_2$) interaction strengths. The patch and nonpatch residue-size parameters are $\sigma_1 = 0.65$ and $\sigma_2 = 0.45$ nm, respectively. The sum of the two contributions gives the ensemble-averaged $R_g$.

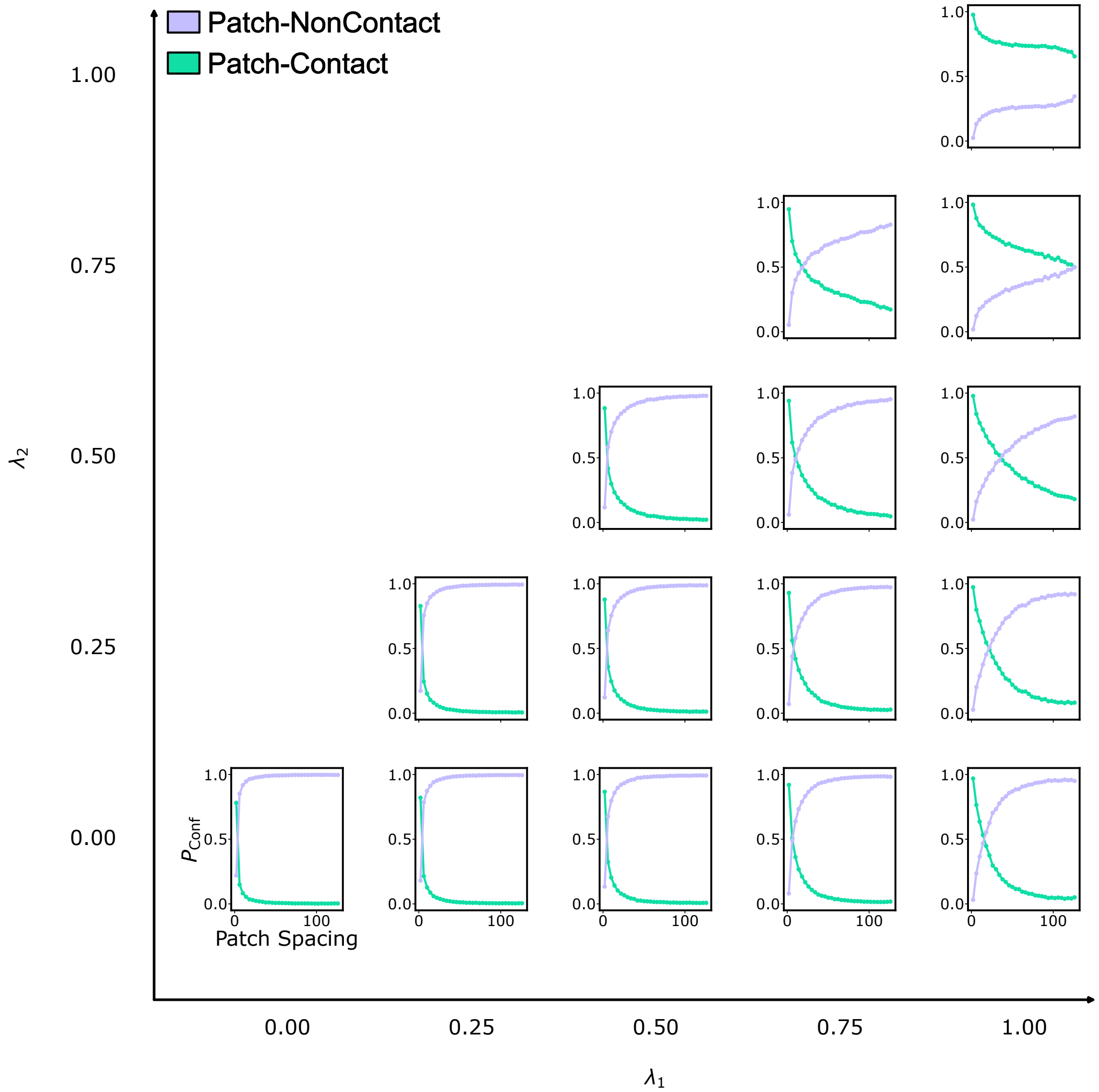


Figure S14: **Conformational-class populations for $\sigma_1 = 0.65$ and $\sigma_2 = 0.45$ nm.** Population fractions of the patch-contact (turquoise) and patch-noncontact (light purple) conformational classes as a function of hydrophobic patch spacing across the sampled combinations of $\lambda_1$ and $\lambda_2$. The patch and nonpatch residue-size parameters are $\sigma_1 = 0.65$ and $\sigma_2 = 0.45$ nm, respectively.

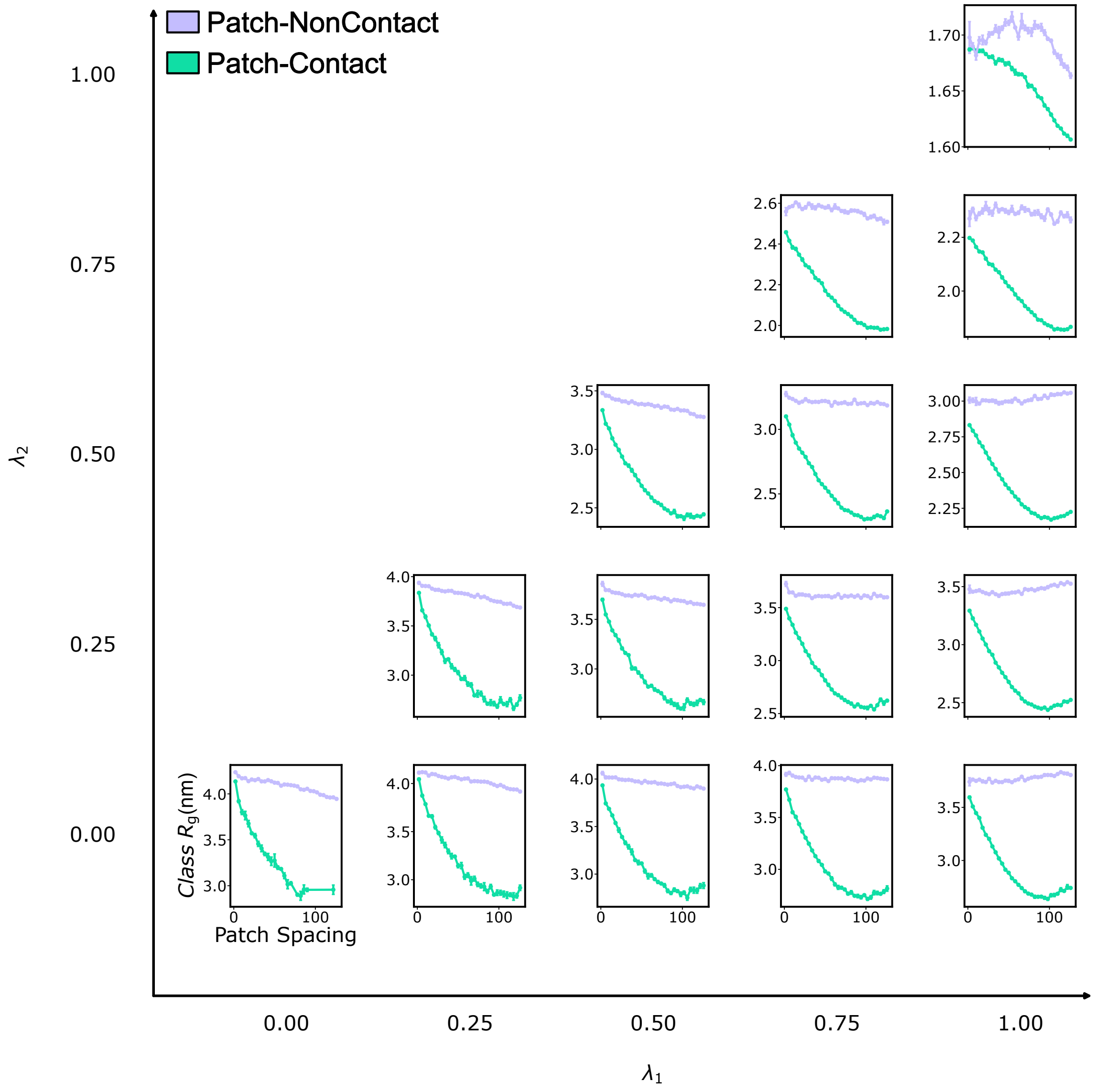


Figure S15: **Class-averaged $R_g$ for $\sigma_1 = 0.65$ and $\sigma_2 = 0.45$ nm.** Class-averaged $R_g$ of the patch-contact (turquoise) and patch-noncontact (light purple) conformational classes as a function of hydrophobic patch spacing across the sampled combinations of $\lambda_1$ and $\lambda_2$. The patch and nonpatch residue-size parameters are $\sigma_1 = 0.65$ and $\sigma_2 = 0.45$ nm, respectively.
